# Comparative genomics and pathogenicity of *Pantoea stewartii* subsp. *stewartii* reveal multiple introductions and limited distribution in Europe

**DOI:** 10.1101/2025.11.07.687269

**Authors:** Aleksander Benčič, Joël F. Pothier, Tanja Dreo

**Author notes:** Corresponding author: Tanja Dreo, Department of Biotechnology and Systems Biology, National Institute of Biology, Vecna pot 121, SI-1000 Ljubljana, Slovenia.

## Abstract

*Pantoea stewartii* subsp. *stewartii* (Pss), the causal agent of Stewart’s wilt in maize, is native to North America but has been detected on several occasions also in Europe, including in Italy in association with seed maize production. These findings have raised concern due to the bacterium’s quarantine status in the EU. To investigate its diversity and possible introduction routes, we performed whole-genome sequencing of Slovenian and other Pss isolates and compared them with publicly available genomes. Comparative genomics revealed that Pss forms a distinct clade within *P. stewartii*, exhibiting high average nucleotide identity (>99.9%). Most of the Slovenian isolates clustered closely together, forming a separate branch consistent with at least two independent introduction events. They were genetically distinct from recent Italian isolates. Additionally, several unique plasmids and prophages were identified in Pss isolates, with notable diversity in mobile genetic elements despite overall genomic homogeneity. Pangenome analysis confirmed an open pangenome for *P. stewartii*, with greater genomic diversity among non-Pss strains. Functional analysis identified multiple secretion systems in Pss, which are likely to contribute to its pathogenicity and insect-mediated transmission. Our findings highlight previously unrecognised diversity within the subspecies and confirm the presence of at least two distinct clades within *P. stewartii*. So far, the detections in Slovenia remain confined to the Vipava Valley with evidence of multiple introduction events. Together, these findings cannot definitely conclude whether Pss is already established in Slovenia. This suggests that eradication remains achievable through continued surveillance and preventive phytosanitary measures.

## INTRODUCTION

*Pantoea stewartii* subsp. *stewartii* (Smith 1898) is a Gram-negative, xylem-inhabiting bacterium and the causal agent of Stewart’s wilt of maize (*Zea mays* L.). The disease, first described in the United States in the late 19^th^ century, is considered one of the most significant bacterial diseases of maize in its native range. Initially classified as part of the genus *Erwinia*, it was later reclassified as part of the genus *Pantoea* (1). Bacteria from this genus inhabit a variety of ecological niches and have been isolated all around the world. *P. stewartii* subsp. *stewartii* (Pss) is native to North America, but findings of the disease have been reported worldwide. Sweet corn and some elite inbred maize lines are especially susceptible to the disease, which has two main phases: wilt and leaf blight. In the first phase, known as the wilt phase, young seedlings become infected, and water-soaked lesions appear on the leaves. The seedlings then wilt severely and often die. The second phase, known as the blight phase, occurs when mature plants become infected and characteristic linear yellow-brown lesions appear, running parallel to the leaf veins. The disease is transmitted from plant to plant via the corn flea beetle (*Chaetocnema pulicaria*), which can also carry it in its gut during the winter months (2). Pss spreads over longer distances via infected seeds, which has led to restrictions on international trade in seed material. This can be mitigated by planting resistant maize cultivars and testing imported seeds for the bacterium’s presence. In addition to maize, Pss can infect other plants, including sudangrass, oats, triticale, sorghum, millet, and sugarcane. Another subspecies, *P. stewartii* subsp. *indologenes* (Psi), infects a variety of crops, including *Allium* spp. and rice (3). Although it is widely accepted that Psi does not cause symptoms of infection in maize, there have been recent reports of Psi isolated from symptomatic maize plants (4).

Pss is an attractive laboratory model organism due to its fast growth rate and the relatively small number of pathogenicity mechanisms it possesses. The two main pathogenicity factors (PF) of Pss are the Hrp type III secretion system (T3SS) and the exopolysaccharide stewartan. The Hrp gene cluster is responsible for the T3SS and the effectors involved in colonising the host plant. Pss does not possess a large number of different effector proteins, indicating that it acquired the T3SS relatively recently (5). One such protein is WtsE, an AvrE-family effector that causes cell death and leads to water-soaking lesions and necrosis in maize plants, playing an essential role in pathogenesis (6). Stewartan, on the other hand, is responsible for vascular streaking, bacterial oozing, and wilting (7). Pss possesses a quorum-sensing (QS) system, which is regulated by *esaI* (a *luxI* homologue) that encodes the signal *N*-3-oxohexanoyl homoserine lactone synthase, and by *esaR* (a *luxR* homologue) that encodes the response regulator (8). The QS system and the Rcs phosphorelay have been observed to regulate the stewartan production, thereby contributing to virulence (9). Another mechanism through which QS is involved in pathogenicity in plants is the regulation of surface motility. Pss exhibits flagella-dependent surface motility, which is involved in biofilm development and plays a significant role in colonising the plant host. The flagellum apparatus is encoded by the *fliTSDC1* locus; mutations in this locus result in loss of motility (10).

The first complete genome sequence of the Pss DC283 strain reveals a 4.53-Mb circular chromosome, ten circular plasmids ranging in size from 4,277 to 304,641 bp, and one linear phage plasmid (ppDSJ01), which is related to the *E. coli* N15 prophage (11). The two smallest plasmids, pDSJ01 and pDSJ02, have medium copy numbers, while the remaining plasmids have low copy numbers (fewer than 10 copies). Genes associated with the T3SS are located on two separate plasmids: pDSJ08 and pDSJ10. The Pss DC283 genome contains genes encoding several putative endoglucanases, xylanases, and a β-1,4–β-1,3 mixed-linkage glucan glucanohydrolase. These enzymes can digest plant cell walls and may play a critical role in enabling the bacteria to access carbohydrates associated with xylem cell wall structure and development (12). The Pss genome also contains many repetitive transposable sequences, which present challenges in genome assembly when only short reads are used. Additionally, multiple prophage sequences were identified within the DC283 genome assembly (11).

In recent years, Pss has been detected on several occasions outside its native range, including in Italy in association with seed maize production and in Slovenia, where findings have so far been confined to the Vipava Valley, a small region with sub-Mediterranean climatic conditions bordering Italy. So far, detections have been limited to symptomatic maize plants appearing late in the growing season, with no evidence of early-season infections, suggesting that *Pss* does not overwinter or cycle naturally under conditions in the areas studied. While previous comparative studies provided valuable insights into the taxonomy of *P. stewartii*, the intraspecific relationships within the species and the genomic diversity among European isolates have remained poorly understood. In particular, it is unclear whether the recent detections represent local establishment, transient survival, or repeated introductions.

The present study aimed to clarify the genetic position and diversity of European Pss isolates and to relate these findings to their phenotypic and pathogenic characteristics. To this end, high-quality whole-genome sequences of Slovenian isolates and additional historical Pss strains were generated and compared with publicly available genomes of *P. stewartii*. Comparative genomic, phylogenetic, and pangenome analyses were combined with pathogenicity testing to assess the diversity and functional potential of Pss strains. The results provide the first detailed genomic and phenotypic characterisation of Pss in Europe, offering new evidence of multiple introduction events, limited distribution, and informing ongoing surveillance and eradication programmes.

## MATERIAL AND METHODS

### Preparation of material

In the scope of this study, 24 new strains of *Pantoea stewartii* subsp. *stewartii* (Pss) were selected for sequencing, including 11 Slovenian isolates and 13 isolates from bacterial strain collections (Table 1). The study includes Slovenian isolates collected between 2018 and 2022. Those were obtained from symptomatic maize plants, which were collected as part of an annual official survey conducted by the Administration of the Republic of Slovenia for Food Safety, Veterinary and Plant Protection, Ministry of Agriculture and Environment, and Phytosanitary Inspectorate. The presence of *P. stewartii* was screened in DNA extracted from symptomatic maize samples using real-time PCR (13). If a sample tested positive, an additional real-time PCR test (14) was performed to differentiate between the subspecies *stewartii* and *indologenes*. The bacterial isolates from maize extracts were identified as Pss by MALDI TOF mass spectrometry (smartfleX, Bruker, Billerica, USA) and real-time PCR tests (14-15). The DNA used for sequencing library preparation was extracted from bacteria that were grown on NA (Bacto Nutrient Agar; Difco) media at 28 °C. Bacterial suspensions were prepared in 10 mM phosphate-buffered saline (PBS; 1.08 g Na_2_HPO_4_, 0.4 g NaH_2_PO_4 ·_ 2H_2_O, 8 g NaCl, 1 L distilled water, pH 7.2). DNA was extracted from the bacterial suspensions using QuickPick™ SML Plant DNA kits (Bio-Nobile, Turku, Finland) and an automated KingFisher™ mL system (Thermo LabSystems), as previously described (15).

**Table 1:** A report on the whole genome sequencing and genome assembly for all *Pantoea stewartii* subsp. *stewartii* strains sequenced within the scope of this study. The table includes metrics on the quality of genome assemblies includes size in megabases (Mb), number of contigs, guanine-cytosine content (GC%), number of coding sequences (CDSs) and number of transfer RNAs (tRNAs), shortest contig length that needs to be included for covering 50% of the genome (N50), completeness as percentage of complete BUSCOs (BUSCO (C) %), contamination percentage as determined by CheckM2 (Cont. %), number of complete plasmids present in genome assembly (Plasmids), and the accession number of genome assemblies in European Nucleotide Archive.

| Strain<sup>a</sup> | Genome size (Mb) | Contigs | GC% | CDSs | tRNAs | N50 | BUSCO (C) % | Cont. % | Plasmids | Acc. No. |
| --- | --- | --- | --- | --- | --- | --- | --- | --- | --- | --- |
| CFBP 1719 | 5.3 | 13 | 53.8 | 5,411 | 74 | 4,480,175 | 98.6 | 0.35 | 8 | GCA_977010205 |
| CFBP 2502 | 5.5 | 15 | 53.7 | 5,694 | 74 | 4,610,069 | 98.5 | 0.45 | 7 | GCA_977010145 |
| CFBP 3157<sup>C</sup> | 5.5 | 15 | 53.6 | 5,762 | 74 | 4,609,570 | 98.6 | 0.59 | 11 | GCA_977011095 |
| CFBP 3165 | 5.1 | 14 | 53.9 | 5,237 | 74 | 4,547,336 | 98.3 | 0.16 | 10 | GCA_977011085 |
| CFBP 3166<sup>C</sup> | 5.4 | 12 | 53.7 | 5,625 | 74 | 4,600,323 | 98.7 | 0.58 | 12 | GCA_977011055 |
| CFBP 3167<sup>T</sup> | 5.2 | 14 | 53.9 | 5,325 | 74 | 4,500,371 | 98.4 | 0.33 | 8 | GCA_977013035 |
| CFBP 3169<sup>C</sup> | 5.1 | 11 | 53.8 | 5,285 | 74 | 4,572,251 | 98.4 | 0.08 | 11 | GCA_977011105 |
| CFBP 3393<sup>C</sup> | 5.3 | 12 | 53.8 | 5,525 | 74 | 4,567,718 | 98.2 | 0.26 | 14 | GCA_977011075 |
| CFBP 3394 | 5.3 | 15 | 53.8 | 5,524 | 74 | 1362,061 | 98.6 | 0.31 | 4 | GCA_977011115 |
| CFBP 3395<sup>C</sup> | 5.3 | 12 | 53.8 | 5,477 | 74 | 4,556,511 | 98.5 | 0.31 | 10 | GCA_977011135 |
| CFBP 3396 | 5.4 | 62 | 53.6 | 5,578 | 70 | 243,803 | 98.6 | 0.71 | 10 | GCA_977011145 |
| CFBP 3445<sup>C</sup> | 5.5 | 15 | 53.7 | 5,730 | 75 | 4,598,621 | 98.7 | 0.58 | 8 | GCA_977011165 |
| NCPBP 3253<sup>C</sup> | 5.5 | 11 | 53.7 | 5,741 | 75 | 4,653,398 | 98.7 | 0.5 | 11 | GCA_977011125 |
| NIB Z 2806<sup>C</sup> | 5.4 | 13 | 53.7 | 5,614 | 75 | 4,579,505 | 98.7 | 0.5 | 14 | GCA_977009295 |
| NIB Z 2809<sup>C</sup> | 5.5 | 13 | 53.7 | 5,687 | 75 | 4,626,859 | 98.7 | 0.5 | 10 | GCA_977010175 |
| NIB Z 3057<sup>C</sup> | 5.4 | 13 | 53.7 | 5,671 | 75 | 4,619,739 | 98.7 | 0.36 | 11 | GCA_977011175 |
| NIB Z 3386<sup>C</sup> | 5.5 | 11 | 53.7 | 5,679 | 75 | 4,626,474 | 98.6 | 0.49 | 10 | GCA_977011155 |
| NIB Z 3388 | 5.5 | 16 | 53.6 | 5,730 | 75 | 4,684,951 | 98.5 | 0.51 | 10 | GCA_977009985 |
| NIB Z 3391 | 5.2 | 203 | 53.7 | 5,441 | 69 | 134,045 | 98.7 | 0.42 | 9 | GCA_977011785 |
| NIB Z 3394 | 5.4 | 55 | 53.7 | 5,644 | 75 | 498,560 | 98.7 | 2.04 | 10 | GCA_977011765 |
| NIB Z 3506 | 5.5 | 17 | 53.7 | 5,678 | 75 | 4,625,939 | 98.6 | 0.58 | 10 | GCA_977011795 |
| NIB Z 3516<sup>C</sup> | 5.4 | 11 | 53.7 | 5,669 | 75 | 4,624,962 | 98.7 | 0.58 | 4 | GCA_977011805 |
| NIB Z 3526 | 5.4 | 14 | 53.7 | 5,655 | 75 | 4,626,258 | 98.7 | 0.57 | 12 | GCA_977011775 |
| NIB Z 3532 | 5.4 | 16 | 53.7 | 5,569 | 75 | 4,558,672 | 98.6 | 0.47 | 12 | GCA_977011755 |

<sup>a</sup> Superscript codes following the strain name correspond to: <sup>C</sup>, complete genomes; <sup>T</sup>, type strain of *Pantoea stewartii* subsp. *stewartii*

### High-throughput DNA sequencing

Whole genome sequencing was performed on DNA extracted from selected bacterial strains (**Table 1**), using the Illumina platform (Illumina, San Diego, USA) alongside one of two long-read sequencing technologies: either PacBio (Pacific Biosciences, Menlo Park, USA) or Nanopore (Oxford Nanopore Technologies, Oxford, UK). Libraries for Illumina sequencing were prepared using a Nextera XT DNA Library Preparation Kit (Illumina) and sequenced on a MiSeq instrument (Illumina) in 2 × 300 bp (V3) mode. Nanopore sequencing was performed using the MinION platform with R9.4.1 or Flongle R9.4.1 flow cells (Oxford Nanopore Technologies). Libraries for Nanopore sequencing were prepared using a Ligation Sequencing Kit (SQK-LSK109) and Native Barcode Expansions 1–12 (EXP-NBD104) and 13–24 (EXP-NBD114), following the manufacturer’s protocol. Magnetic beads (Mag-Bind TotalPure NGS, Omega Bio-Tek, Norcross, USA) were used to purify the DNA during library preparation. Demultiplexing and basecalling were carried out using the Guppy basecaller v6.4.2. PacBio sequencing was performed by an external provider (Novogene GmbH, Munich, Germany). HiFi SMRTbell libraries were prepared and sequenced for PacBio sequencing using the PacBio Sequel II system (Pacific Biosciences).

### Bioinformatic analysis

The basecalled nanopore reads were trimmed using Porechop v0.2.4 (16) to remove adapters, and then filtered using NanoFilt v2.8.0 (17) to remove low-quality reads. The PacBio raw reads were used to generate circular consensus sequences with the pbccs v6.4.0 package (Pacific Biosciences), which were then used for genome assembly in subsequent analyses. Hybrid genome assembly using short and long reads was performed with Unicycler v0.4.8 (18), which uses SPAdes v3.15.5 (19) to assemble contigs from short reads. These contigs were then used alongside long reads to assemble hybrid genomes using miniasm v0.3-r179 (20) and Racon v1.4.20 (21). To assemble genomes using only long reads, Flye assembler v2.8.1-b1676 (22) was used. Assemblies were manually curated to remove any duplicated contigs. If possible, gaps in hybrid assemblies were filled with sequences generated using only long read assemblies. BUSCO v5.7.1 (23) with the pantoea_odb12 dataset was used to determine the completeness of genomes. Completeness was additionally determined by CheckM2 v1.1.0 with database v3, which was also used to determine contaminations within genomes (24). The assembled genomes were visualised and aligned using the commercially available software Geneious Prime v2024.0.7 and the Mauve plugin v1.1.3 (25). Automatic annotation of the assembled genomes was performed using Bakta v1.9.3 and the v5.1 database (26). Genome assemblies of newly sequenced strains were deposited in the European Nucleotide Archive under the study PRJEB100601.

### Phylogenetic analysis

The phylogenetic analysis used 24 newly sequenced Pss strains (Table 1), 38 published genome assemblies from different strains in the NCBI GenBank database, and three genomes assembled from raw data from the Sequence Read Archive (Table 2). All genomes were assembled using the same protocol described in the chapter ‘Bioinformatic analysis’. The genomes included in Table 1 and Table 2 are referred to simply as ‘genomes’ through the text. Genomes with incomplete metadata (host, location, and year of isolation), poor quality (high fragmentation, BUSCO (C) score under 95% and low sequencing depth), duplicate genomes of the same strain, and metagenome-assembled genomes were excluded from the analysis.

**Table 2:**
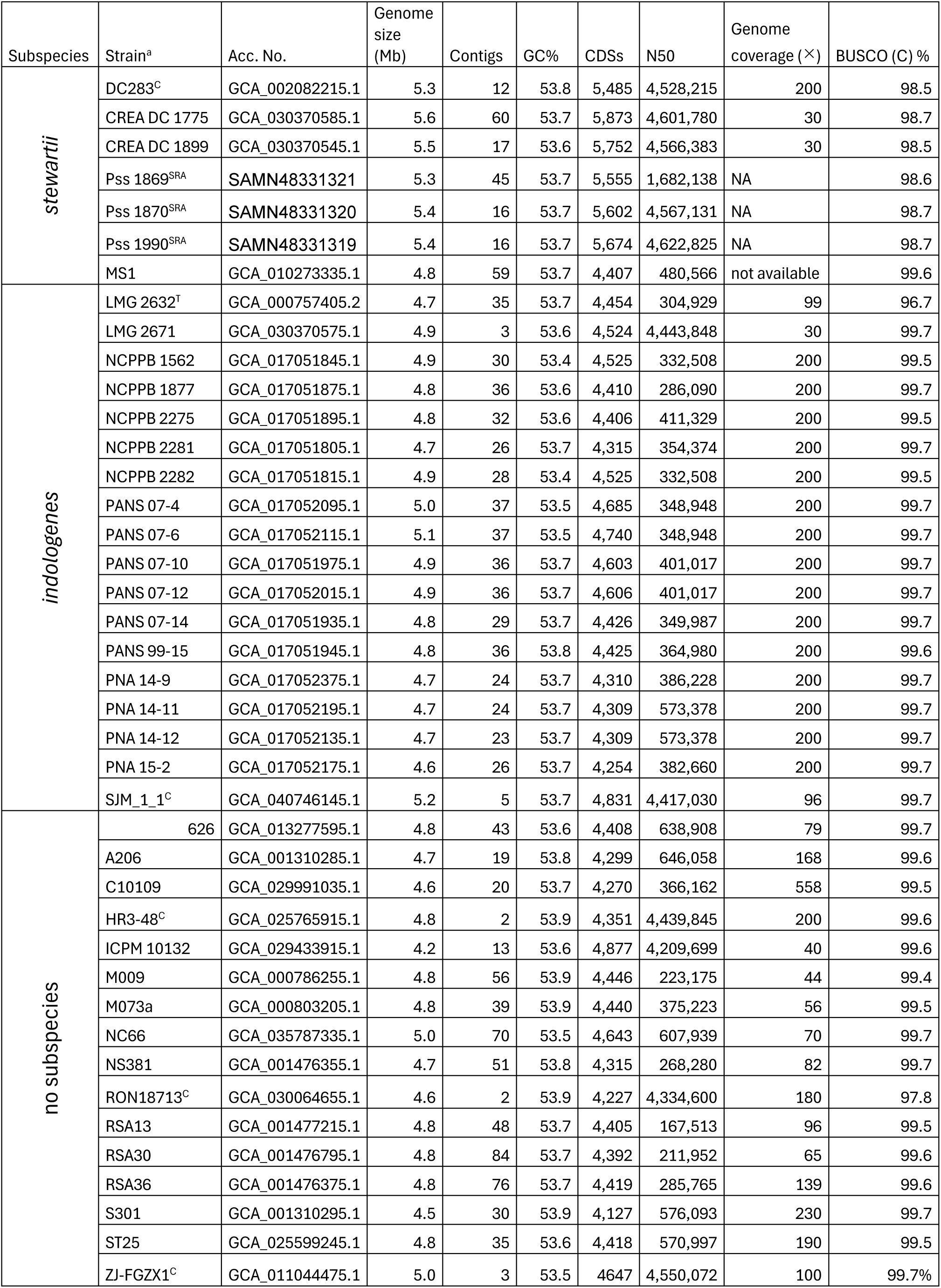
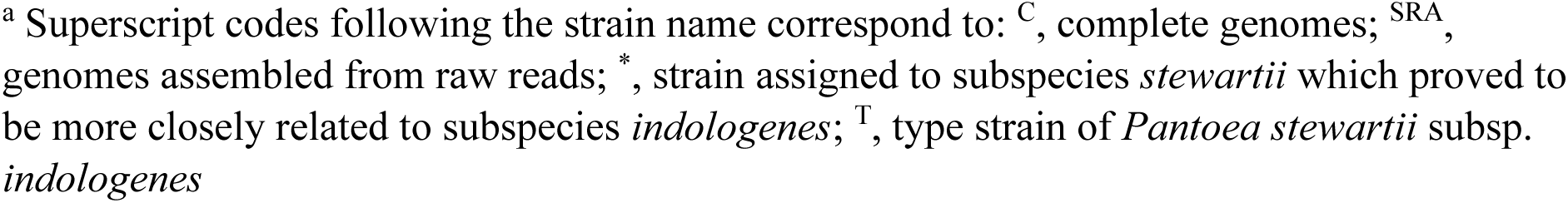
The publicly available *Pantoea stewartii* genomes used in this study are listed below. Data on genome assemblies includes: GenBank or Sequence Read Archive accession number, size in megabases (Mb), number of contigs, guanine-cytosine content (GC%), number of coding sequences (CDSs), shortest contig length that needs to be included for covering 50% of the genome (N50), genome coverage (×) as provided in NCBI database and completeness as percentage of complete BUSCOs (BUSCO (C) %). Additional information on genomes assembled from raw sequences is available in Supplement Table S3.

Within the NCBI database, the genomes were categorised as either subspecies *stewartii* or *indologenes,* or neither. For the phylogenetic analysis of all *P. stewartii* genomes, average nucleotide identity (ANI) was used. This analysis was performed using the bioinformatic tool fastANI v1.33 (27) to generate an ANI matrix. A phylogenetic tree was constructed from the ANI matrix with the bioinformatic tool SplitsTree App v6.5.1 (28) using the neighbour-joining method. For the phylogenetic analysis of genomes belonging to the Pss, single-nucleotide polymorphisms (SNPs) were utilised. The programme snippy v4.6.0 (29) was employed to identify SNPs between the haploid reference genome DC283 and other Pss genomes, and to perform a core genome alignment. The maximum likelihood criterion and the programme RaxML-NG v0.9.0 were used to construct the phylogenetic tree(30). Phylogenetic analysis of subspecies Pss was additionally performed using the accessory genome (genes present in less than 95 % of analysed genomes) as well. Presence-absence genes output generated in analysis performed by software Roary v3.13.0, described in chapter ‘Pangenome analysis’, was used to generate a binary matrix, which was then used to calculate the Jaccard distance matrix using the R package *vegan* v2.7-2 (31). A phylogenetic tree was constructed from a distance matrix using neighbour-joining method with the R package *ape* v 5.8-1 (32). Phylogenetic networks were calculated from both core genome SNPs alignment and accessory genome binary matrix in SplitsTree App using *p*-distances and neighbour net method. To perform whole genome multi-locus strain typing (MLST) of Pss genomes, the software chewBBACA v3.4.1 (33) was used together with a training file created from genome DC283 by software Pyrodigal v3.6.3 (34, 35). A whole genome MLST scheme was generated and used to perform allele calling. Core genome MLST was determined after removing paralogs and the allele call on the number of loci present in all genomes was used to generate a minimum spanning tree which was generated in R v4.0.5 and the integrated development environment RStudio v1.4.1106 (36), using packages ape (32), tidyverse v2.0.0 (37), ggnetwork v0.5.14 (38) and igraph v 2.1.4 (39) as previously described (40).

### Pangenome analysis

The pangenomes of the subspecies Pss and Psi, as well as the species *P. stewartii*, were determined using annotated.gff3 files generated by Bakta, and the Roary pipeline was used to calculate the pangenome (41). The analysis included the newly sequenced genomes of Pss and *P. stewartii* genomes from NCBI databases (Table 1, Table 2). The results were visualised using the programming language R v4.0.5 (42), in combination with the ggplot2 v3.3.3 (43), ggpubr v0.4.0.999 (44) and gridExtra v2.3 (45) packages.

### Pathogenicity factors and mobile genetic elements

A literature survey was conducted to identify known and potential pathogenicity factors for Pss in maize (Supplement Table: S1). Sequences of proteins involved in the type III secretion system were extracted from the annotation of the published Pss DC283 genome (GCF_002082215.1). A BLASTp analysis was conducted on protein sequences generated in the annotation of genomes to identify the presence of different types of secretion system components and other pathogenicity factors. Pathogenicity factors were considered present if they were identified with a score of over 80% identity and query coverage. A secretion system was considered present if over 60% of the proteins that form it were identified. Prophage-like regions and genes in *P. stewartii* genomes were identified using the PHASTEST v3.0 web server (PHAge Search Tool with Enhanced Sequence Translation: https://phastest.ca/) (46). To determine the identity of plasmids in complete genome assemblies, a BLASTn analysis was performed on contigs and the Core Nucleotide BLAST database. To identify plasmids within fragmented genome assemblies at the contig level, BLASTn was used to identify sequences of known plasmids within the analysed assemblies. The plasmids were first extracted from the complete genome of DC283, which served as a reference. Rearrangements in the plasmids were analysed using progressiveMauve v1.1.3 (25) and visualised using the R package genoPlotR v0.8.11 (47).

### Pathogenicity test on plants

A pathogenicity test was performed on the sweetcorn variety ‘Gucio’ (L’Ortolano S.r.l., Cesena, Italy) using the following *P. stewartii* strains: CFBP 1719, CFBP 3165, CFBP 3167^T^, CFBP 3169, CFBP 3445, CFBP 3614^T^, NIB Z 2806, and NIB Z 3391. Each group contained ten plants, with strains CFBP 1719 and NIB Z 2806 duplicated in two groups (giving a total of 20 plants per group). The strains used in the pathogenicity test were blinded to ensure unbiased reporting of symptoms. Maize seedlings inoculated with 10 mM PBS served as the negative control, while the Pss strain NIB Z 2806 served as the positive control. The maize seeds were sterilised in 1.5% sodium hypochlorite solution (v/v) before being sown into 0.46 L pots containing a special substrate (Hawita Gruppe GmbH, Vechta, Germany). The maize seedlings were then grown in controlled greenhouse chambers following a 16 h light (22 °C)/8 h dark (19 °C) day/night cycle and 60% humidity. Fourteen days after sowing, the maize seedlings in growth stages V2-3 were infected via stem inoculation under the first leaf using a sterile 25 G hypodermic needle. This was performed using a bacterial suspension of 10⁸ cells/mL (as determined by optical density/McFarland measurements at a wavelength of 565 nm), prepared from an approximately 48-hour culture on King’s B medium. Following inoculation, the plants were incubated at a constant temperature of 26 °C, with 85% humidity and under the same lighting conditions as before inoculation. The presence of Pss infection symptoms was examined in the plants at 3-, 7-, 14-, and 30-days post-inoculation (DPI). The severity of the symptoms was evaluated using a scale from 0 to 7: 0 represented no symptoms; 1, yellow stripes on the leaves; 2, necrotic lesions on the leaves; 3, small, localised, water-soaked lesions with ooze; 4, larger, water-soaked lesions with ooze; 5, leaf deformities with widespread lesions; 6, stunted growth with wilting; and 7, plant death. The scale was modified from that described by Mohammadi et al. (12). Photos of representative symptoms used for symptom evaluation are presented in Supplement: Table S2.

## RESULTS

### Genome sequencing of Pss strains

A total of 24 Pss strains were newly sequenced, comprising 11 Slovenian isolates and 13 strains from bacterial collections. All strains were sequenced using Illumina technology, with either ONT or PacBio technology used to obtain hybrid assemblies. The genome assembly sizes of the newly sequenced Pss strains ranged from 5.1 Mb (CFBP 3165) to 5.5 Mb (CFBP 3157), with an average size of 5.4 Mb. For strains for which a complete chromosome was obtained, the size ranged from 4.49 Mb (CFBP 1719) to 4.68 Mb (NIB Z 3388), with no major rearrangements observed (Table 1, Supplement: Figure S1, Table S3). These results are consistent with data from the publicly available assembly of strain DC283, which has a genome size of 5.2 Mb and a chromosome size of 4.53 Mb. Differences in genome size are mainly due to differences in plasmids present in different strains. The number of coding DNA sequences (CDSs) ranged from 5,237 (CFBP 3165) to 5,944 (CFBP 1719), averaging 5,604. The GC% of Pss ranged from 53.6% to 53.9%, averaging 53.7% (Table 1, Supplement: Table S3). When compared to other genome assemblies assigned to *P. stewartii* in GenBank, Pss genomes had, on average larger size and a higher number of CDSs. Genome sizes of other *P. stewartii* strains ranged from 4.2 Mb and 4,016 CDSs (RON18713) to 5.2Mb and 4,736 (ICMP 10132), with an average genome size being 4.8 Mb and an average number of CDSs 4,359 (Table 2). All newly sequenced and assembled genomes had high completeness (over 98%) and low contamination (under 3 %) and could be included in further analysis (Table 1, Supplement: Table S3).

### Phylogenetic analysis

#### *Species* P. stewartii

The genomes of *P. stewartii* included in the phylogenetic analysis had previously been assigned to either the Pss or Psi subspecies, or to no subspecies at all. To obtain reliable results, only the complete genome assemblies and high-quality draft assemblies with BUSCO completeness score (C) > 95 % were included in phylogenetic and other analyses. These genomes were used to calculate an ANI matrix between the genomes, which was then used to construct a phylogenetic tree. The analysis revealed that the *P. stewartii* phylogenetic tree formed two distinct branches. Strains assigned to Pss formed a single monophyletic group (Figure 1). The second branch included all strains assigned to the Psi subspecies, as well as all unassigned strains, all of which were more closely related to Psi. The exception was strain MS1, which was isolated from jackfruit and was previously assigned to Pss but is genetically more closely related to Psi strains. In the following text, all 35 strains from the branch containing Psi strains will be referred to as Psi. Strains in the Psi branch showed much greater diversity, with ANI values among them ranging from 98.48 % to 99.99% and a mean of 98.92%. In contrast, Pss strains were found to be very closely related, with ANI values ranging from 99.7% to 99.99% and a mean of 99.93%. ANI percentages between strains in Pss in Psi clades ranged from 98.27% to 98.76% with an average ANI being 98.56% (Supplement Table S3). Results of phylogenetic analysis based on ANI clearly demonstrate Pss and Psi to form two distinct lineages within species *P. stewartii*.

**Figure 1:**
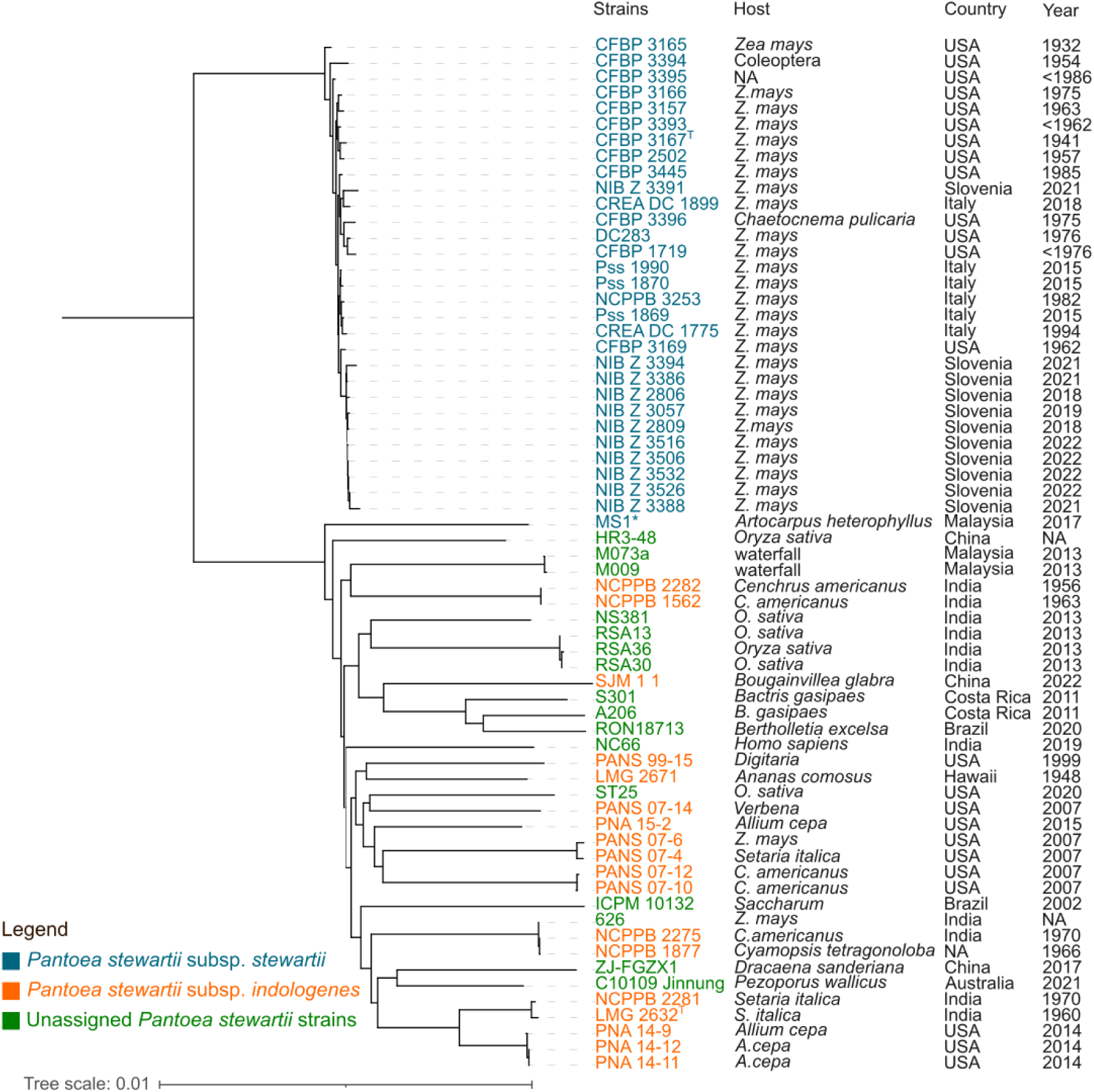
Phylogeny of the *Pantoea stewartii* species. A phylogenetic tree based on the average nucleotide identity (ANI) of 65 *P. stewartii* genomes, obtained from NCBI GenBank or assembled in this study. Strains belonging to the *stewartii* (Pss) subspecies are shown in cyan, those belonging to the subspecies *indologenes* (Psi) in orange, and strains not assigned to any subspecies in green. The figure also shows metadata including the geographic origin (country), year of isolation, and host organism. All Pss strains form a separate branch within the phylogenetic while all strains without an assigned subspecies are dispersed among the Psi strains within the phylogenetic tree. ^T^: strains CFBP 3167^T^ and LMG 2632^T^ are type strains of subspecies *stewartii* and *indologenes*, respectively. *: This indicates strain MS1, which was previously assigned to subspecies *stewartii,* but which is more closely related to other *P. stewartii* strains.

#### *Subspecies* stewartii

The Pss branch comprised 30 strains, including 11 Slovenian isolates, six Italian isolates, and the remainder from the USA. To investigate the relationships between these closely related strains, phylogenetic analysis of Pss subspecies was performed on both core and accessory genomes. In core genome-based phylogeny, SNPs were identified within the core genome alignment, which was then used to construct a phylogenetic tree using the maximum likelihood method. The results showed that all Slovenian isolates, except for NIB Z 3391, form a single branch within the Pss branch (Figure 2: A, Supplement: Figure S3). Slovenian isolates were most closely related to the Pss branch containing strains CFBP 3396, CFBP 3166, CFBP 3157, and CREA DC 1899. In accessory genome-based phylogeny binary matrix of genes present in the accessory genome was generated and used to construct a phylogenetic tree using Jaccard distances and the neighbour-joining method. Results of accessory genome-based phylogeny showed similar patterns. One major difference is that the Slovenian strain NIB Z 3388 does not group with the majority of other Slovenian isolates but with another Slovenian isolate, NIB Z 3391, which is positioned separately from other Slovenian isolates (Figure 3: A). Phylogenetic network analysis showed the core genome produces results with fewer conflicts in comparison to the accessory genome. It additionally showed strain NIB Z 3388 to form a separate branch (Figure 2: B, Figure 3: B). The Italian isolates do not group with the Slovenian isolates in any of the performed phylogenetic analyses, but they exhibit a similar pattern of behaviour. Five of them form a distinct branch, while isolate CREA DC 1899 is positioned elsewhere within the Pss tree (Figure 2, Supplement: Figure S3). To further examine the relationships between Pss strains, cgMLST was performed, which used a minimum spanning tree (MST) to visualise those relationships. The number of allelic differences between nodes in MST ranges from 4 to 90, which indicates that Pss strains do not form a unified clonal complex. Slovenian strains, with the exceptions of NIB Z 3388 and NIB Z 3391, showed a smaller number of allelic differences among them (4 to 13) in comparison to the other strains (number of allelic differences over 18). A similar pattern is again seen with Italian strains, which further form a separate group except for strain CREA DC 1899 (Figure 2: C).

**Figure 2:**
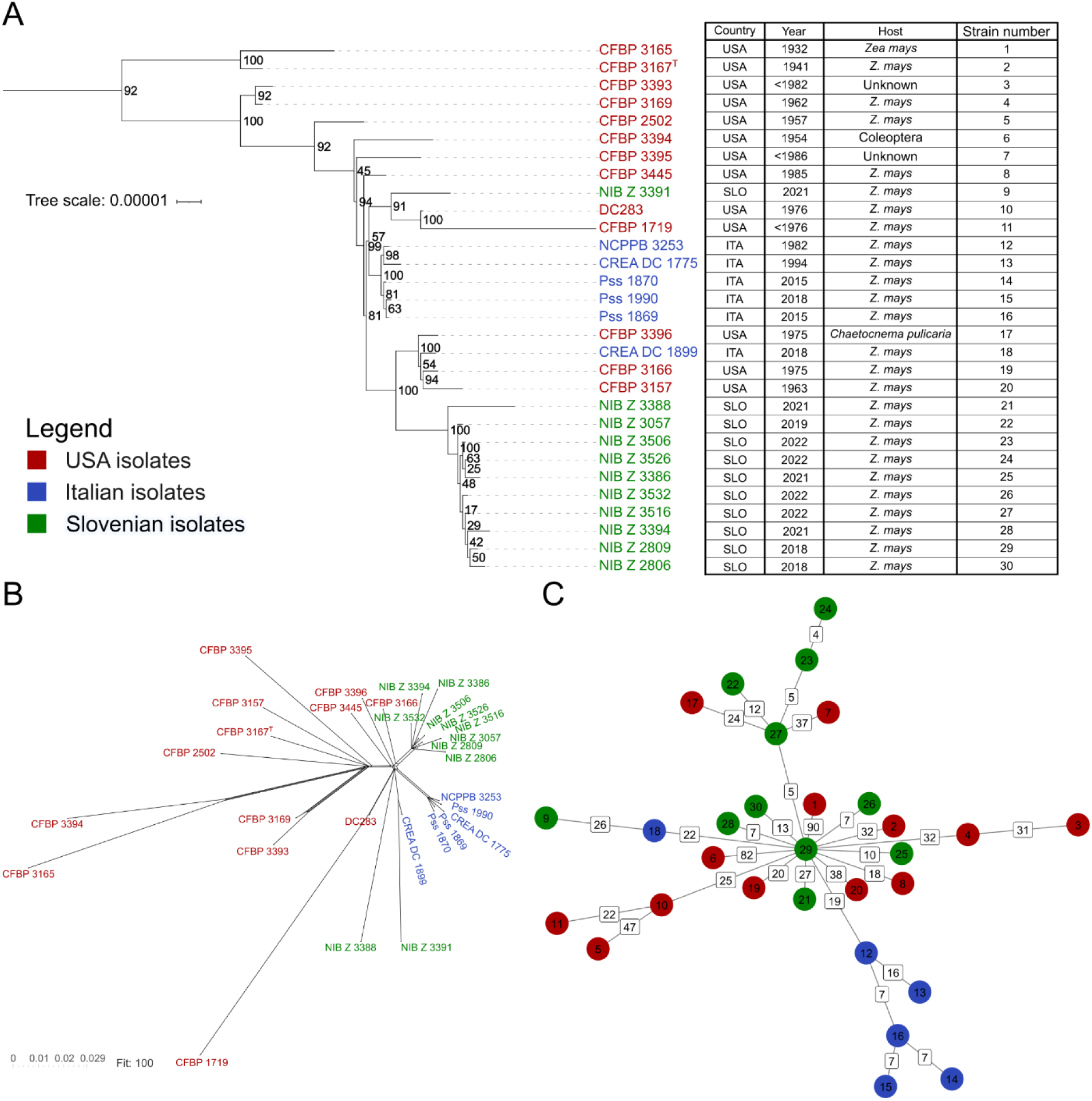
Core genome-based phylogenetic analysis of *Pantoea stewartii* subsp. *stewartii*. **A**: Phylogenetic tree constructed from high-quality *P. stewartii* subsp. *stewartii* (Pss) genomes single nucleotide polymorphisms alignment and maximum likelihood method with 1000 bootstrap replicates. The bootstrap values are shown as percentages in the nodes of the tree. The figure further shows metadata, including the geographic origin (country), year of isolation, and host organism strain numbers. **B**: Phylogenetic network of Pss constructed using P distances. **C:** Minimum spanning tree generated from core genome multiple locus strain typing showing allelic differences between strains. Node contains strain numbers of strains as listed in the table in panel A. Names of the strains are coloured according to country of origin in the whole figure, with Slovenian strains shown in green, the USA strains in red, and the Italian strains in blue as shown in the legend.

**Figure 3:**
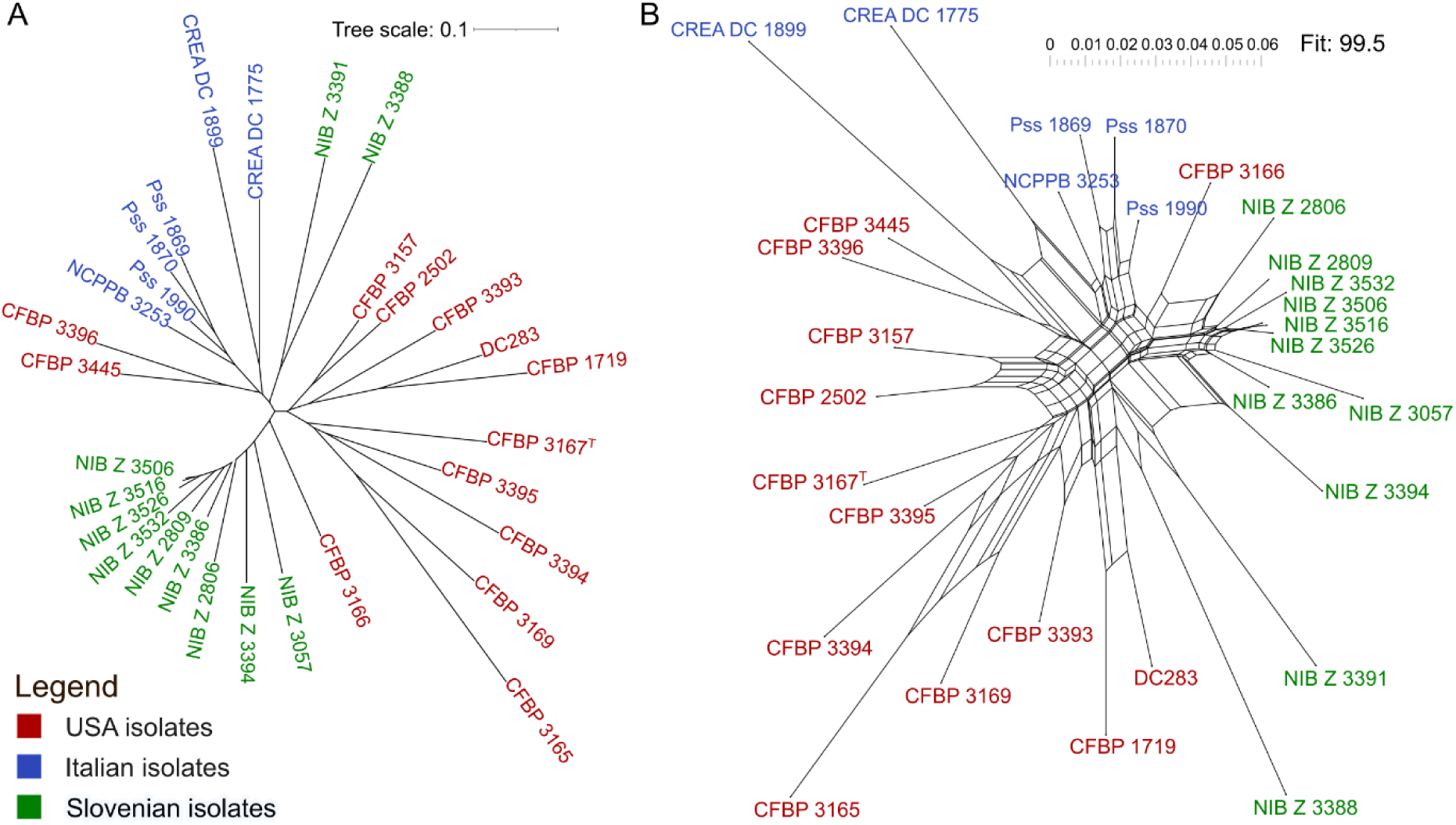
Accessory genome-based phylogenetic analysis of *Pantoea stewartii* subsp. *stewartii*. **A**: Unrooted phylogenetic tree of *P. stewartii* subsp. *stewartii* (Pss) constructed from the accessory genome (genes present in under 95% genomes) binary matrix using Jaccard distances and neighbour-joining clustering method. **B**: Phylogenetic network of Pss constructed using *p* distances. Names of the strains are coloured according to country of origin in the whole figure, with Slovenian strains shown in green, the USA strains in red, and the Italian strains in blue, as shown in the legend.

### Pangenome

The pangenome was determined for both subspecies, as well as for the entire *P. stewartii* species. The complete pangenome of Pss contained 7,425 genes. Of these, 3,968 formed the core genome, while the extended soft core — genes present in at least 95% of genomes — consisted of an additional 443 genes. The shell contained an additional 1,584 genes present in 15% to 95% of the genomes, and the cloud contained a further 1,430 genes present in less than 15% of the genomes (Figure 4 Figure 4A, B; Table 3). The Psi clade pangenome consisted of 11,952 genes, 3,496 of which were core genes (Figure 4Figure 4 C, D), whereas the pangenome of all *P. stewarti*i species consisted of 15,559 genes, of which 3,013 were core genes (Figure 4Figure 4 E, F; Table 3). Core genes represented the largest percentage of the pangenome for subspecies Pss (53.4%), whereas in Psi and the entire *P. stewartii* species, core genes represented a smaller percentage of the pangenome (29.3 and 19.4%, respectively; Figure 4**, G**). The number of genes in the pangenome continues to increase with additional genomes added, indicating *P. stewartii* possesses open pangenomes (Figure 4 A, C, E).

**Figure 4:**
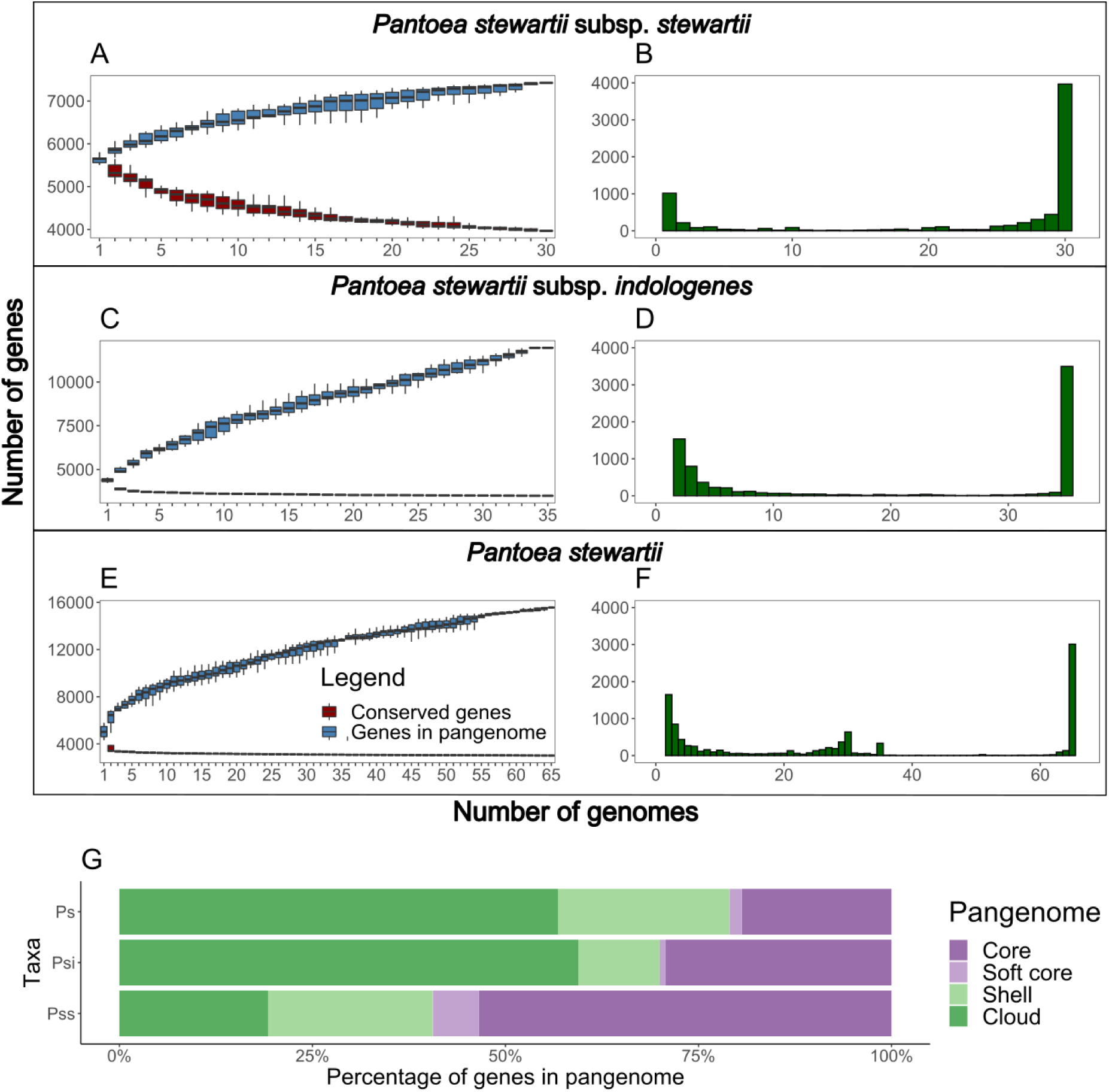
Pangenome analysis of *Pantoea stewartii* and its subspecies. The pangenomes of Pantoea stewartii subsp. stewartii (Pss), strains of *P. stewartii* subsp. *indologenes* (Psi), and the whole *P. stewartii* species are shown. Panels A, C, and E illustrate how the pangenome’s gene count increases with each additional genome (in blue), while the core genome’s count decreases with each additional strain, stabilising at a set number of core genes characteristic of *P. stewartii*. Panels B, D, and F show the number of genes present in a given number of genomes. Panels A and B show the data for Pss, panels C and D show the data for non-Pss strains, and panels E and F show the data for all *P. stewartii* strains. Panel G shows the pangenome composition for Pss, Psi, and *P. stewartii*, expressed as a percentage of genes included in the core, soft core, shell, or cloud.

**Table 3:** Pangenome composition for *Pantoea stewartii* subspecies *stewartii* (Pss), subspecies *indologenes* (Psi), and the whole species *P. stewartii*. The table includes the criteria for inclusion and number of genes in core, soft core, shell, cloud and the total number of genes for each group.

| Pangenome | Criteria | Pss ($N = 30$) | Psi ($N = 35$) | *P. stewartii* ($N = 65$) |
| --- | --- | --- | --- | --- |
| Core genes | From 99 to 100% strains | 3,968 | 3,496 | 3,013 |
| Soft core genes | From 95 to 100% strains | 443 | 92 | 250 |
| Shell genes | From 15 to 95% strains | 1,584 | 1,260 | 3,456 |
| Cloud genes | From 0 to 15% strains | 1,430 | 7,104 | 8,840 |
| Total genes | All strains | 7,425 | 11,952 | 15,559 |

### Diversity of mobile genetic elements

#### Plasmid diversity

The genomes forming the Pss branch of the phylogenetic tree were analysed for the presence of plasmids. The analysis identified 15 different plasmids, one of which was linear (ppPSS01) and the rest circular (Supplement Table S5). Despite being closely related, the analysed Pss strains showed diversity in their plasmids, with the number of plasmids ranging from eight (CFBP 3167^T^) to 14 (CFBP 3445 and CFBP 3157), and the majority of genomes contained more than 10 plasmids (Figure 5), with no strain containing all plasmids. The larger plasmids pPSS10 (64.9 kb), pPSS11 (72.0 kb), pPSS13 (107.8 kb), and pPSS14 (305.8 kb) were present in some form in all analysed genomes (Figure 5, Supplement Table S6). Two previously unidentified plasmids, pPSS06 and pPSS09, were discovered in newly sequenced Pss strains. Plasmid pPSS06 is 36.3 kb in size on average, has a GC% of 43.3, and is present in nine genomes, while pPSS09 is 54.5 kb in size on average, has a GC% of 48.8, and is present in 20 genomes (Figure 5, Supplement Table S5-6). Both new plasmids contain a type IV secretion system (T4SS), including VirB10/TraB/TrbI, TrbG/VirB9, and TrbC/VirB2. They also contain a P-type DNA transfer ATPase, VirB11, which is involved in horizontal gene transfer via conjugation. Some rearrangements of plasmids have been observed in different Pss strains compared to the DC283 reference genome.

**Figure 5:**
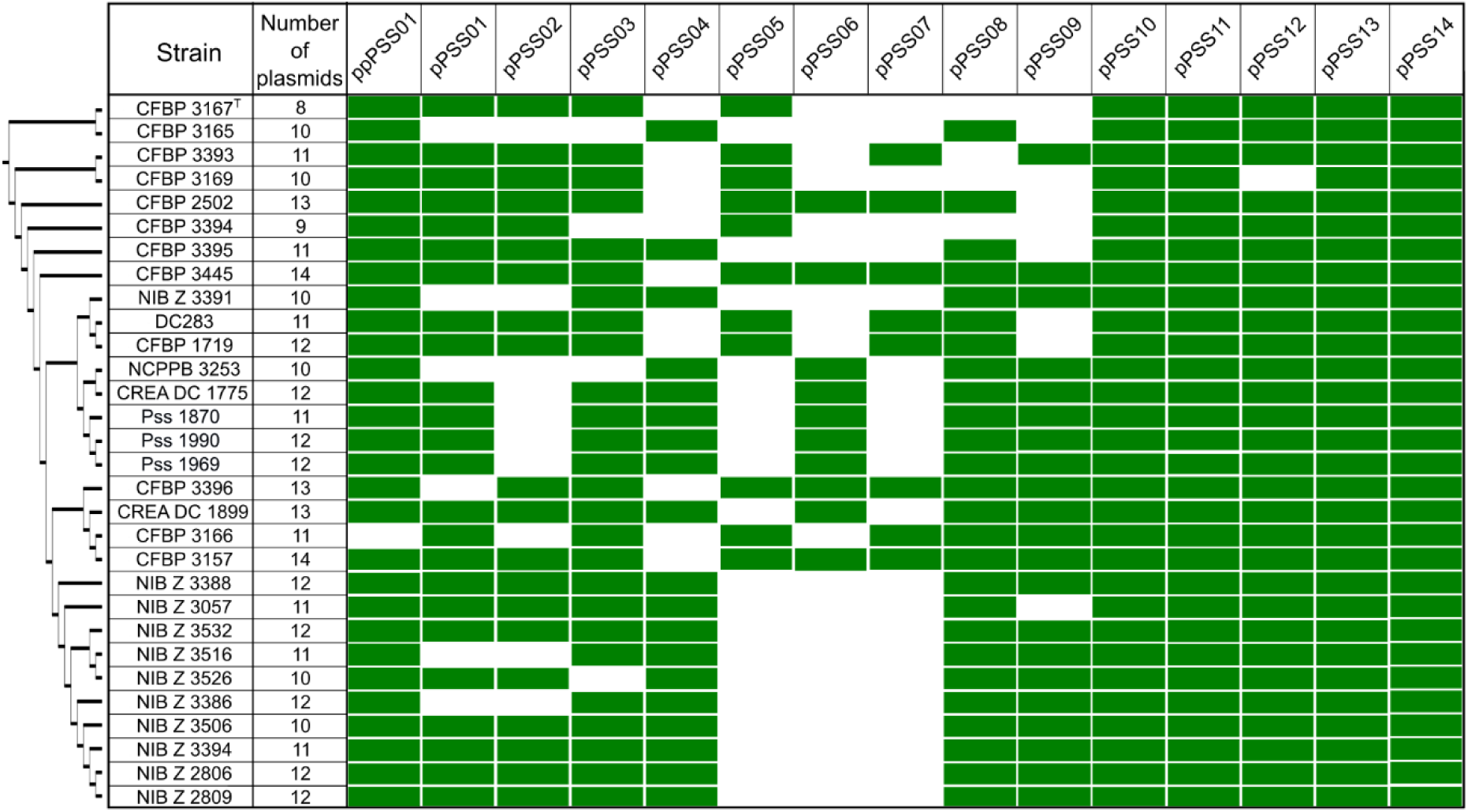
Plasmid diversity of *Pantoea stewartii* subsp. *stewartii*. Strains are listed according to their position within the core genome phylogenetic tree, which is presented as a dendrogram on the left side of the figure. The presence of plasmids in strains is indicated by a green colour, while an empty field indicates that the plasmid was not identified in these genome assemblies. A total of 15 different plasmids were identified among the 30 *Pantoea stewartii* subsp. *stewartii* (Pss) strains. The number of plasmids per strain ranges from eight (CFBP 3167^T^) to 14 (CFBP 3445), and no strain contains all known plasmids. Details about the plasmids and the nomenclature used in this figure can be found in Supplemental Table S4.

Additionally, multiple arrangements were also observed between plasmids in different strains. Most of these rearrangements were observed in the 133 kb plasmid pDSJ09 from reference genome DC283, which is split into the plasmids pPSS11 and pPSS12 in all Pss genomes except said reference genome and CFBP 1719. This is in agreement with the phylogenetic analysis, in which those two strains are positioned next to each other. The pPSS12 plasmid contains an insertion absent from pDSJ09. The exception is strain CFBP 3169, which lacks pPSS12 altogether. The size of both pPSS11 and pPSS12 is just over 70 kb in most Pss genomes, with some notable exceptions. Strain CFBP 3165 has a smaller pPSS11 plasmid (45 kb). pPSS12 showed greater variability, ranging in size from 52 to 91 kb (Supplement: Table S6, Figure S2). Other plasmids also showed some rearrangements and variability. The plasmid pPSS14 is conserved among all Pss strains but is smaller in strains CFBP 3165 (151 kb) and CFBP 3169 (144 kb) than in all other strains, in which pPSS14 is over 300 kb in size (Supplement: Figure S3). In 19 strains, parts of plasmids pPSS05 and pPSS07 merged to form plasmid pPSS04; however, large parts of both original plasmids were lost. This variant is present in all Slovenian and Italian isolates but absent from the majority of American isolates (Supplement: Table S6 and Figure S4). In 14 strains, plasmid pPSS08 has a large insertion that forms a monophyletic branch in the phylogenetic tree (Supplement: Table S6 and Figure S5). This variant of pPSS08 is present in all Slovenian isolates except NIB Z 3391, which is consistent with the ANI-based phylogenetic analysis. Some rearrangements were observed in individual strains only. In strain CFBP 3157, part of plasmid pPSS10 was inserted into plasmid pDSJ03, resulting in a smaller pPSS10 (47 kb) and a larger pPSS03 (34 kb) (Supplement: Figure S6). In strain CFBP 3394, part of plasmid pPSS10 was inserted into pPSS05, resulting in smaller pPSS10 (48 kb) and larger pPSS10 (45 kb) (Supplement: Figure S7). Overall, Slovenian strains exhibit a consistent plasmid composition among them, with eight plasmids present in all analysed genomes. Similar can also be said for Italian insoles, while American isolates, on the other hand, show greater diversity in plasmid presence. Of all the plasmids identified in Pss genomes, only pPSS14 is also present in other *P. stewartii* strains. The pPSS14 plasmid was identified in all analysed strains, but it is smaller in Psi strains, which lack the part encoding the T3SS.

#### Prophages

All the *P. stewartii* genomes used in the phylogenetic analysis were also examined for the presence of prophages. This analysis revealed that Pss strains have a greater number of prophage regions and genes than Psi strains. Pss genomes contain between 10 and 14 intact prophage regions, ranging in size from 10.5 to 112.6 kb (Supplement Table S7-9). In addition to complete prophages, all genomes contained additional incomplete prophage regions. The genomes of the Pss strains exhibited a distinct prophage profile, with the prophages Burkho_BcepMu_NC_005882, Entero_phiT5282H_NC_049429, Haemop_SuMu_NC_019455, and Salmon_SEN34_NC_028699 being conserved across all of them (see Supplementary Table S8). One prophage was found to be in the form of the linear plasmid ppDSJ01, present in 29 out of 30 analysed Pss genomes. Genomes in the Psi branch contained between 0 and 6 complete prophage regions, ranging in size from 10.5 to 70.1 kb. Only one prophage, Erwini_ENT90_NC_019932, was conserved in the majority of strains; however, it is in Pss present only in genomes of CFBP 1719 and Pss 1990 (Supplement Table S8-9).

### Pathogenicity factors and secretion systems

The two main pathogenicity mechanisms in Pss are the type III secretion system (T3SS) and the production of the exopolysaccharide stewartan. However, Pss contains many other factors that contribute to its virulence in maize. To determine the presence of different pathogenicity factors (PF) in Pss and other *P. stewartii* strains, the annotations of the 24 genomes sequenced in the present study and the 41 genomes from the NCBI databases were analysed (Table 1, Table 2).

#### Secretion systems

The Pss genome contains three different T3SS operons: one is encoded by the pPSS14 plasmid (T3SS-1), and the other two are encoded by the pPSS13 plasmid (T3SS-2 and T3SS-3). All Pss genomes contain at least two T3SSs; 27 genomes contain all three. T3SS-1, which is required for Pss pathogenicity in maize, is absent from the genomes of CFBP 3165 and CFBP 3169. T3SS-2, which is linked to bacterial colonisation of the corn flea beetle gut, is present in all analysed genomes. The function of T3SS-3 is not fully understood, but it is present in all Pss genomes except the type strain CFBP 3167^T^ (Figure 6). The majority (*n* = 28) of strains in the Psi branch also contain some type of T3SS. T3SS-1 is present in 24 Psi genomes, while the other two T3SSs found in Pss are absent from the genomes of other *P. stewartii* strains. Strains NCPPB 1562, NCPPB 2282, SJM_1_1, and ST25 possess a different T3SS each, indicating different acquisition pathways. NCPPB 1562 and NCPPB 2282 have the same type of T3SS; however, the T3SSs of SJM_1_1 and ST25 are unrelated. Strains MS1, NRRL B-13, NS381, RSA13, RSA30, RSA36, and S301 lack the complete T3SS apparatus and the associated effectors (Figure 6).

**Figure 6:**
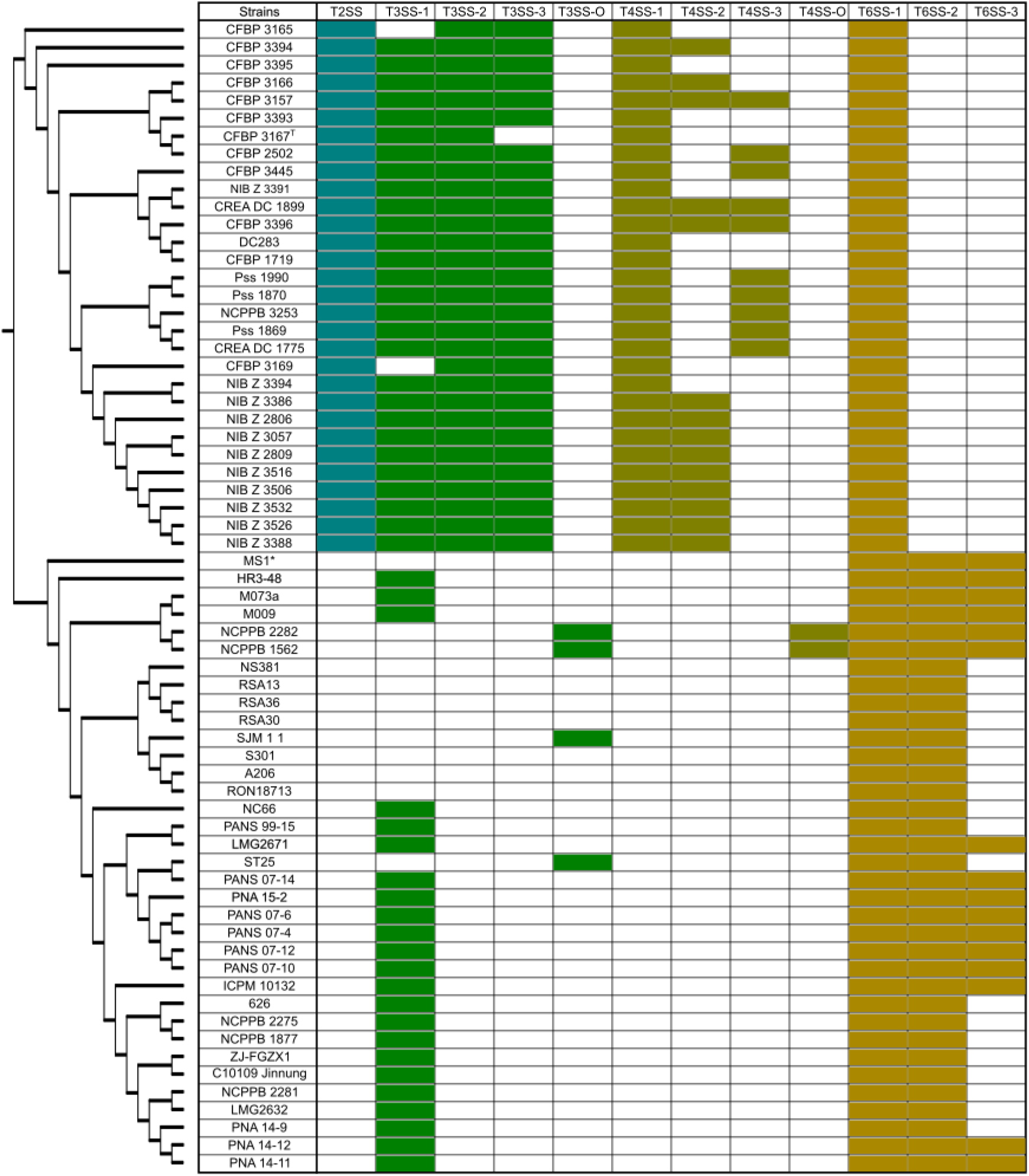
Diversity of secretion systems among *Pantoea stewartii* strains. This figure shows the presence or absence of the following secretion systems, as identified using the genome annotations of 65 *Pantoea stewartii* strains: type II (T2SS; turquoise), type III (T3SS-1, –2, –3, –O; green), type IV (T4SS-1, –2, –3, –O; olive green), and type VI (T6SS-1, –2, –3; ochre). A white field indicates the absence of a secretion system in this strain. The strains are listed in order of their position in the dendrogram shown on the left.

In addition to the T3SS, *P. stewartii* strains contain other types of secretion systems. A type II secretion system (T2SS) is encoded by plasmid pPSS13, which is present in all analysed Pss genomes, but absent in all Psi branch strains. All Pss genomes contain a type IV secretion system (T4SS), which is encoded by plasmids absent in other *P. stewartii* strains. The genomes of strains NCPPB 1562 and NCPPB 2282 contain a T4SS that differs from that of Pss strains. Pss genomes containing the newly discovered plasmids pPSS06 and pPSS09 also contain an additional T4SS involved in horizontal gene transfer by conjugation. All *P. stewartii* genomes contain nonfunctional parts of a type VI secretion system (T6SS) encoded on the bacterial chromosome. Unlike Pss t Psi strains also encode for at least one additional T6SS locus, which is thought to play an important role in the plant pathogenicity of these strains. Nineteen genomes encode an additional third T6SS locus (Figure 6).

#### Other pathogenicity factors

In addition to secretion systems, *P. stewartii* strains also have many other genes involved in pathogenicity that have been reported in the literature. A survey was conducted to identify 54 known and possible PFs that were, at the time of analysis, described in the existing literature. These PFs were used in further analysis (Supplement Table S1). These include genes involved in bacterial adhesion and aggregation, capsule formation, carbohydrate degradation, carotenoid biosynthesis, flagellum-mediated surface motility, the inducible oxidative stress response, transport across the bacterial membrane, quorum sensing, the secretion of effector proteins, the acquisition of iron via siderophores, and the biosynthesis of stewartan and water soaking during infection. All PFs were identified by BLAST analysis in at least one strain. The majority of the pathogenicity factors were conserved in the analysed Pss strains, with the genes involved in quorum sensing and stewartan biosynthesis being conserved in all analysed *P. stewartii* genomes. All Pss strains also contain cellulase (RefSeq: WP_006122111.1), which, in addition to T2SS, could play a role in pathogenicity. Additionally, gene *ucp1,* located on plasmid pPSS10, which enables gut colonisation in the insect vector, is present in all Pss genomes except Pss 1869; however, it is absent in all Psi genomes. Conversely, *P. stewartii* strains lacking T3SS-1 also lack the associated effectors WtsE and the HrpN family hypersensitivity reaction elicitor. The *wtsE* gene was also not found in the genome of CREA DC 1775, which otherwise contains T3SS-1 and *hrpN*; however, this may also be due to gaps in the genome assembly (Supplement Table S10).

### Pathogenicity test

A pathogenicity test (PT) was performed to determine the pathogenicity of different Pss strains on maize and to study the effects of the different PFs present in these strains. The strains used in the PT were selected to include those lacking different T3SSs, as well as American and Slovenian isolates. The PT also included a Psi type strain, which was expected to cause no symptoms. Plants in the positive control group exhibited symptoms indicative of Pss infection, including yellow stripes on leaves, necrotic water-soaked lesions on leaves, and leaf deformities. In contrast, plants in the negative control group exhibited only yellow stripes near the inoculation site in some plants, resulting from mechanical damage caused by the needle and buffer. Symptoms were most severe in maize seedlings inoculated with the Pss strains CFBP 1719, CFBP 3445, NIB Z 2806, and NIB Z 3391, with an average symptom intensity greater than three at 3, 7, and 14 days post-inoculation (DPI) (Figure 7, Supplement Table S11). These strains were also the only ones still showing symptoms at 30 DPI. The CFBP 3167^T^ strain, which lacks T3SS-3, exhibited milder symptoms that did not result in stunted growth or leaf deformities. These symptoms then subsided, with no symptoms observed at 30 DPI. Pss strains CFBP 3165 and CFBP 3169, which lack T3SS-1, did not exhibit symptoms indicative of Pss infection (Figure 7, Supplement Table S11). The Psi strain CFBP 3614^T^, which contains T3SS-1 but lacks other T3SSs, exhibited minimal symptoms following inoculation (averaging 1.4), which disappeared by 14 DPI and were not indicative of Pss infection (Figure 7, Supplement Table S11). Overall, showed differences in symptom severity between strains, with some strains showing severe symptoms while others lacked symptoms of infection with Pss.

**Figure 7:**
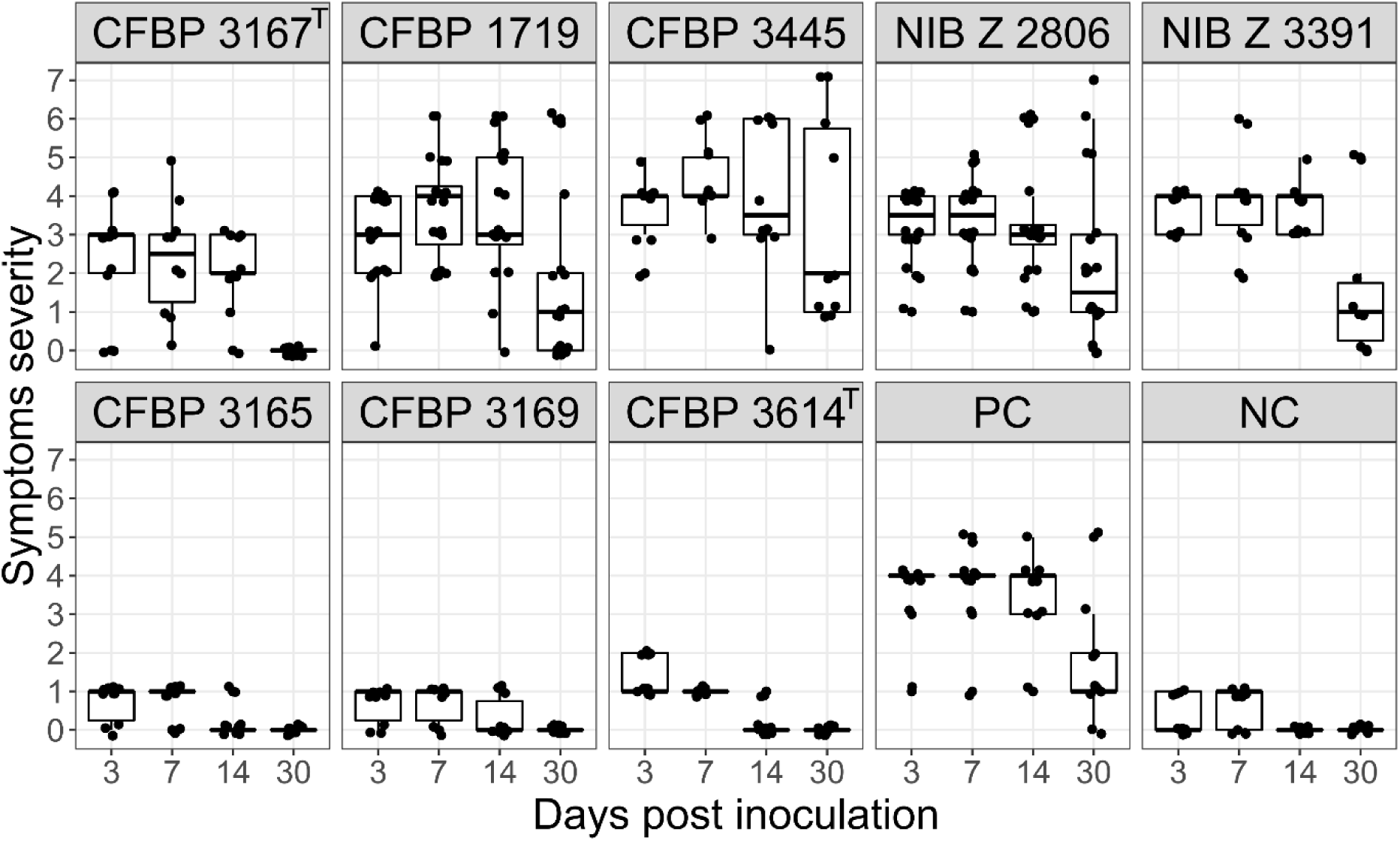
Results of pathogenicity tests on maize for different *Pantoea stewartii* strains. The severity of the symptoms caused by the different *Pantoea stewartii* strains was assessed at 3-, 7-, 14-, and 30-day post-inoculation. Maize seedlings in the negative control (NC) group were inoculated with 10 mM phosphate-buffered saline (PBS), while those in the positive control (PC) group were inoculated with strain NIB Z 2806.

## DISCUSSION

Although *Pantoea stewartii* subsp. *stewartii* (Pss) is native to North America; detections have been reported on other continents as well. Measures have been implemented to prevent its introduction into the European Union, including testing of imported seed lots. Despite these measures, several new detections have been reported in Europe in recent years. Notably, *Pss* has been identified in Italy in association with seed maize production (in years 2022-2024), representing one of the first confirmed occurrences of the pathogen in European commercial seed systems (48, 49). These detections underscore the potential for inadvertent movement of the bacterium through seed trade and highlight the importance of systematic surveillance.

In Slovenia, multiple findings have also been reported in recent years, concentrated within the Vipava Valley in the west of the country, bordering Italy. This region has a distinct sub-Mediterranean climate, which differs from the continental conditions prevailing across most of Slovenia. The genomes of Slovenian isolates and additional Pss strains from bacterial collections were sequenced in the scope of this study, covering detections reported up to 2022. Subsequent findings and isolates are currently under analysis and will be presented separately. These sequences, together with genome data available in public databases, were used to perform phylogenetic and comparative genomic analyses to clarify the position of Slovenian isolates within Pss subspecies, and the position of Pss within the *P. stewartii* species.

Previous comparative genomic studies have shown *P. stewartii* to be a distinct bacterial species, most closely related to *P. ananatis* and *P. allii* (50–52). However, the position of Pss within *P. stewartii* has remained unclear due to the scarcity of high-quality genomic data. At the time of this work, the genome of strain DC283 was the only complete, high-quality assembly on which much of our knowledge of Pss genomics was based, whereas other available assemblies were fragmented or of low completeness. This study generated new, high-quality genomic data that enabled robust comparative analyses of the Pss subspecies and of the *P. stewartii* species as a whole. Phylogenetic analysis based on ANI revealed that *P. stewartii* strains form two distinct clades: one comprising all Pss strains, which display a close relatedness (>99.9% ANI), and another containing all *P. stewartii* subsp. *indologenes* (Psi) and unassigned strains. The latter clade is more diverse, consistent with its broader range of host plants, whereas Pss isolates are largely restricted to maize. Strain MS1, previously assigned to the Pss, was nested within the Psi clade, which supports the findings of a previous study (53).

Given the close relatedness among Pss strains, core-genome SNPs were used for higher-resolution phylogenetic analysis. Slovenian isolates formed a distinct branch in the phylogenetic tree, except for the strain NIB Z 3391, which clustered separately. This pattern suggests at least two independent introduction events, one of which may have resulted in limited local dissemination within Slovenia. Strain NIB Z 3388 may represent an additional introduction, as it shows lower relatedness to the remaining Slovenian isolates and forms a separate branch in both accessory-genome and network analyses. The Italian isolates display a comparable pattern, with most forming a separate branch except for strain CREA DC 1899. The currently available genomes of European Pss isolates show no clear genetic connection between findings from different countries, indicating that these are likely to represent multiple, independent introduction events. Broader surveys and additional genome sequencing from different geographical areas and time periods will be required to clarify the pathways of entry and potential persistence of Pss in Europe.

Despite being closely related, Pss strains displayed diversity in plasmid content. The analysed genomes contained between eight (CFBP 3167^T^) and 14 (CFBP 3445 and CFBP 3157) plasmids, which is higher than in other *P. stewartii* strains. Two previously undescribed plasmids, pPSS06 and pPSS09, were identified. The total plasmid count may still be underestimated in some assemblies, particularly those with low average genome coverage and those assembled using only long reads (49). For example, the publicly available genome of strain ATCC 8199 (GCF_045159535.1), which was not included in the analysis, contains only five plasmids, whereas our assembly of strain CFBP 3167^T^, which represents the same isolate, contains eight. It is improbable that several plasmids were lost during cultivation, especially as one of them, pPSS14, is highly conserved among *Pantoea* species. Genome assemblies, however, remain models of genome organisation, and some apparent rearrangements may not reflect biological variation. Among the plasmids identified, only the pPSS14 occurs in other *Pantoea* species, where it is known as the large *Pantoea* plasmid (LPP-1) and plays an important role in pathogenesis (54). Slovenian isolates share a similar plasmid composition among them, except for isolate NIB Z 3391, consistent with their phylogenetic grouping. Italian isolates show a comparable pattern, again with the exception of CREA DC 1899. The functional importance of plasmids is underscored by the pathogenicity factors they encode: the T2SS on pPSS13; T3SS on pPSS11 and pPSS14; and T4SS on pPSS06 and pPSS09. The presence of these systems likely contributes to the observed diversity of plasmids within Pss. In addition to plasmids, other mobile genetic elements, notably prophages, are abundant in Pss genomes. They contain more prophage regions than those of Psi, suggesting an important evolutionary role of prophages in the diversification of the Pss lineage.

The phylogenetic patterns are mirrored in the pangenome analysis. Pss has a smaller pangenome than Psi, with a higher proportion of core genes, reflecting its lower overall diversity. The species *P. stewartii* has an open pangenome, consistent with its extensive complement of mobile genetic elements and its capacity for horizontal gene transfer (51). The slower increase in gene number within the Pss pangenome likely reflects its narrower host range. Genes unique to Pss may be associated with its pathogenicity in maize and its capacity for insect-mediated transmission.

The presence and composition of pathogenicity factors vary between *P. stewartii* subspecies. The two principal virulence determinants of Pss are the Hrp type III secretion system (T3SS-1) and the exopolysaccharide stewartan (8, 55). The *wce-I*, *-II,* and *-III* gene clusters for stewartan biosynthesis were present in all analysed *P. stewartii*, confirming the potential for exopolysaccharide production. Pss also contains two additional T3SSs: one implicated in colonisation of the corn flea beetle gut, and another of as yet unclear function (56). The limited repertoire of effector proteins supports the view that Pss acquired the T3SS relatively recently (55). One such effector is WtsE, a member of the AvrE family of effectors that causes cell death, resulting in water-soaked lesions and necrosis in maize plants, and is essential for pathogenesis (6). Pss also possesses a T2SS absent from Psi, whereas Psi retains functional T6SS loci that are inactive in Pss (57). Only T3SS-1 is shared by both subspecies, being located on pPSS14, homologous to LPP-1. This indicates an older evolutionary origin of T3SS-1, although its loss in some strains of both subspecies points to relatively recent separate evolutionary events. Previous work showed that *P. stewartii* produces endoglucanase, which cleaves cellulose into smaller subunits and contributes to mobility in xylem vessels and to symptom development (12). Our analysis indicates that the T3SS-3 locus also encodes a cellulase and the expansin protein YoaJ, which binds cellulose and may enhance plant colonisation (58). This could explain why strains lacking T3SS-3 cause milder symptoms that do not progress to systemic infection. However, the further studies are needed to confirm the functional roles of T3SS-3 and YoaJ. Notably, T3SS-3 is absent only from the type strain CFBP 3167^T^, one of the oldest isolates, making it difficult to determine whether this system was lost from that strain or gained in others.

Pathogenicity tests on maize seedlings confirmed the functional significance of the secretion systems. Strains lacking T3SS-1 did not cause typical Pss symptoms, confirming its essential role in virulence on maize. A Psi strain carrying T3SS-1 caused only mild symptoms, supporting the idea that other factors are also required for pathogenicity. The Pss type strain (CFBP 3167^T^), lacking T3SS-3, produced only transient symptoms, further confirming that this system contributes to disease development. This further confirmed its role in the pathogenicity of Pss in maize. Together, these results corroborate the hypothesis that the absence of specific secretion systems markedly reduces virulence.

Pss remains an important maize pathogen, and its recent detection in new regions has renewed attention to its biology and movement. This study helps fill a long-standing knowledge gap by providing complete and high-quality genome assemblies, thereby enabling the placement of new isolates within the broader context of *P. stewartii* diversity. The results offer new insights into the genetics and biology of Pss, including the diversity of its mobile genetic elements and virulence factors. These findings are critical for improving diagnostic specificity and informing measures to limit further introductions. However, more research is required to understand the mechanisms governing its persistence and apparent geographical dispersal. The increasing number of detections in Europe remains difficult to interpret without coordinated monitoring across countries.

The detection of genetically similar isolates in geographically proximate locations may indicate limited local dissemination, but further data are required to confirm this. These findings nonetheless emphasise the need for continued surveillance and phytosanitary vigilance. The situation in Slovenia most likely reflects proactive monitoring rather than a unique occurrence, suggesting that similar introductions may be going undetected elsewhere. Preliminary phenotypic analyses using BIOLOG plates indicate that Slovenian isolates possess broad metabolic versatility consistent with survival under varying environmental conditions, supporting the view that these findings reflect persistence rather than active establishment.

Taking together the available evidence, we cannot yet conclude that Pss is already established in Slovenia. In addition, the bacterium has been detected in a weed species within the affected area, while limited surveys have so far not confirmed the presence of potential insect vectors. The principal North American vector, the corn flea beetle (*Chaetocnema pulicaria*), is not known to be present in Europe and has not been reported from Slovenia. The absence of early-season (spring) infections and the late appearance of symptoms, typically from late summer to autumn, further support the view that active overwintering and the presence of effective vector-mediated transmission have not yet been showcased. Ongoing studies are addressing the potential role of alternative insects and the ecology of Pss in non-crop hosts. These findings collectively indicate that eradication within the affected area remains a realistic objective, provided that targeted phytosanitary measures and continued monitoring are maintained. Strengthening genomic, phenotypic, and ecological surveillance in other regions will therefore be crucial to assess the true frequency and extent of Pss introductions in Europe. Slovenia provides a valuable case study for exploring the genomic diversity of recently detected European isolates. Future efforts will concentrate on refining diagnostic and isolation methods, expanding phenotypic characterisation, and supporting coordinated surveillance to ensure early detection and containment of any further introductions.

## ACKNOWLEDGMENTS

The authors acknowledge the financial support of the Slovenian Research and Innovation Agency (research core funding No. P4-0165, Research Programme on Biotechnology and Systems Biology, and young researchers programme – MR Aleksander Benčič, CRP V4-2415 MapQuest); the Ministry of Agriculture, Forestry and Food of the Republic of Slovenia (Professional Tasks Programme in the Field of Plant Health; Euphresco projects 2018-A275 and 2023-A454; and CRP V4-2415 MapQuest); and the European Union Reference Laboratories – Bacteriology co-funded by the Ministry of Agriculture, Forestry and Food of the Republic of Slovenia; and the Department of Life Sciences and Facility Management of the Zurich University of Applied Sciences (ZHAW), Wädenswil, Switzerland.

We are grateful to Primož Pajk (Administration of the Republic of Slovenia for Food Safety, Veterinary and Plant Protection) for his continuous support and constructive discussions, and to colleagues from the Chamber of Agriculture and Forestry – Institute of Nova Gorica (KGZ-NG) and other regional services for conducting field inspections, visual assessments, and sample collections within the official survey programmes.

We would like to thank S. Baeyen, J. Venneman and J. Van Vaerenbergh from Flanders Research Institute for Agriculture, Fisheries and Food for their help in the implementation of new bioinformatic analyses.

We thank the laboratory staff of the National Institute of Biology, in particular A. Blatnik, V. Dukić, J. Matičič, L. Matičič, M. Pirc, Š. Prijatelj Novak and N. Turnšek, for their valuable technical assistance. The successful isolation of the strains described in this study would not have been possible without their routine yet essential work. We also thank A. Bogožalec Košir for comments on the manuscript.

## Abbreviations

ANI: Average Nucleotide Identity
CDS: Coding DNA Sequence
DPI: Days Post-Inoculation
EPS: Exopolysaccharide
LPP-1: Large *Pantoea* Plasmid 1
MALDI-TOF: Matrix-Assisted Laser Desorption/Ionization Time-of-Flight
NA: Nutrient Agar
PCR: Polymerase Chain Reaction
PBS: Phosphate Buffered Saline
PF: Pathogeny Factor
PT: Pathogenicity Test
QS: Quorum Sensing
rRNA: Ribosomal Ribonucleic Acid
SNP: Single Nucleotide Polymorphism
T2SS: Type II Secretion System
T3SS: Type III Secretion System
T4SS: Type IV Secretion System
T6SS: Type VI Secretion System
tRNA: Transfer Ribonucleic Acid

