## Supplement Figures for "Comparative genomics and pathogenicity of *Pantoea stewartii* subsp. *stewartii* reveal multiple introductions and limited distribution in Europe"

*Figure S1: Alignment showing all complete chromosomes from assembled genomes of* Pantoea stewartii *subsp.* stewartii *strains. Alignment was created using progressiveMauve to identify genome segments that form the backbone of the alignment. Shades of red indicate the same direction of DNA sequence, while shades of blue indicate reversed direction. Strains are ordered based on their position in the phylogenetic tree shown in Figure 2A.*

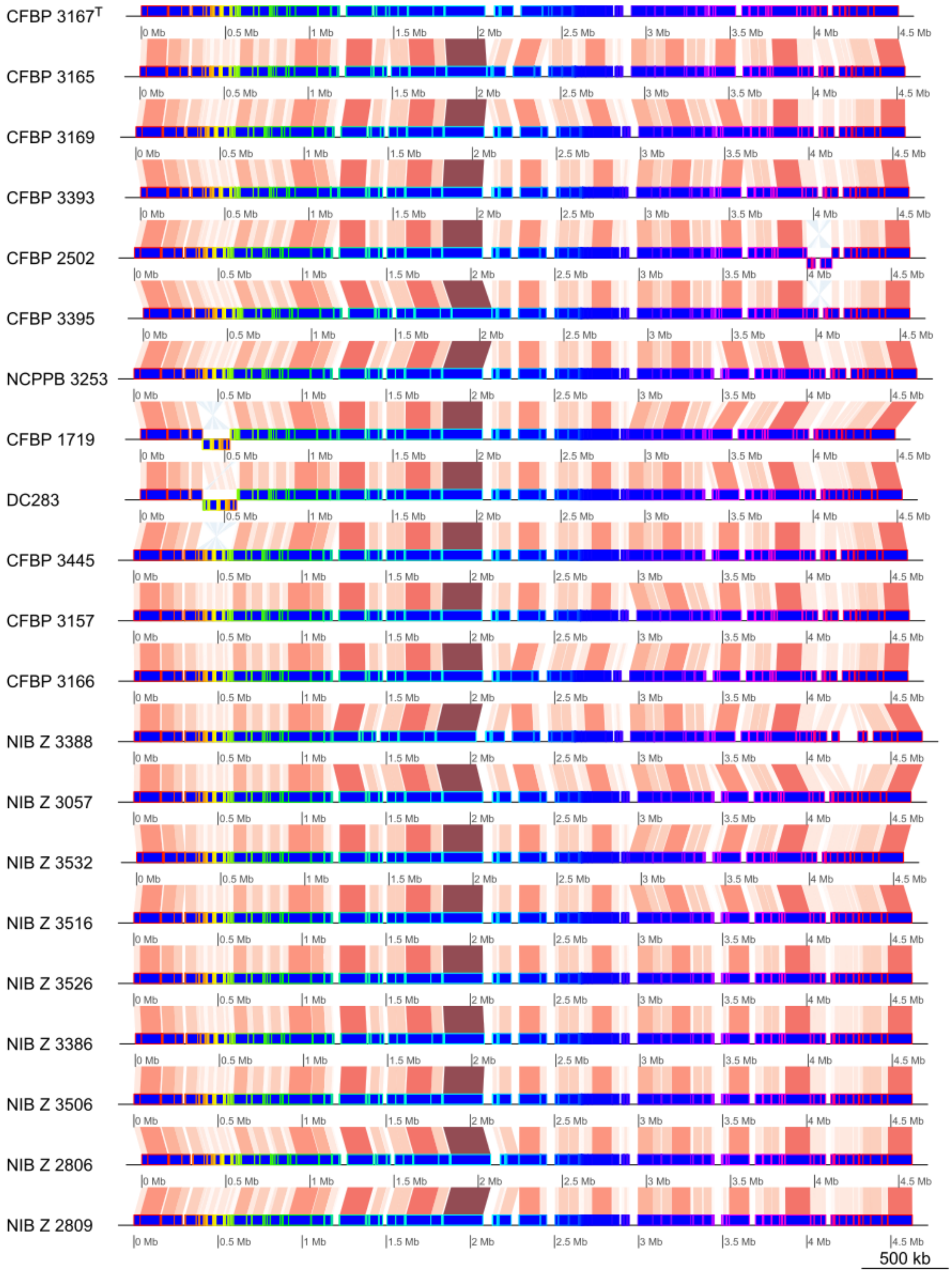

*Figure S2: Alignment showing all complete plasmids pPSS11 and pPSS12 from assembled genomes of* Pantoea stewartii *subsp.* stewartii *strains in which both plasmids are present in complete form. Both plasmids are a bit over 70 kb in size in the vast majority of genomes. The end of pPSS11 and start of pPSS12 is indicated by a dark red line. In the genomes of strains DC283 and CFBP 1719, these two plasmids are joined to form a single plasmid, pDSJ09, which is 133 kb in size. Strains are ordered based on their position in the phylogenetic tree in Figure 2A.*

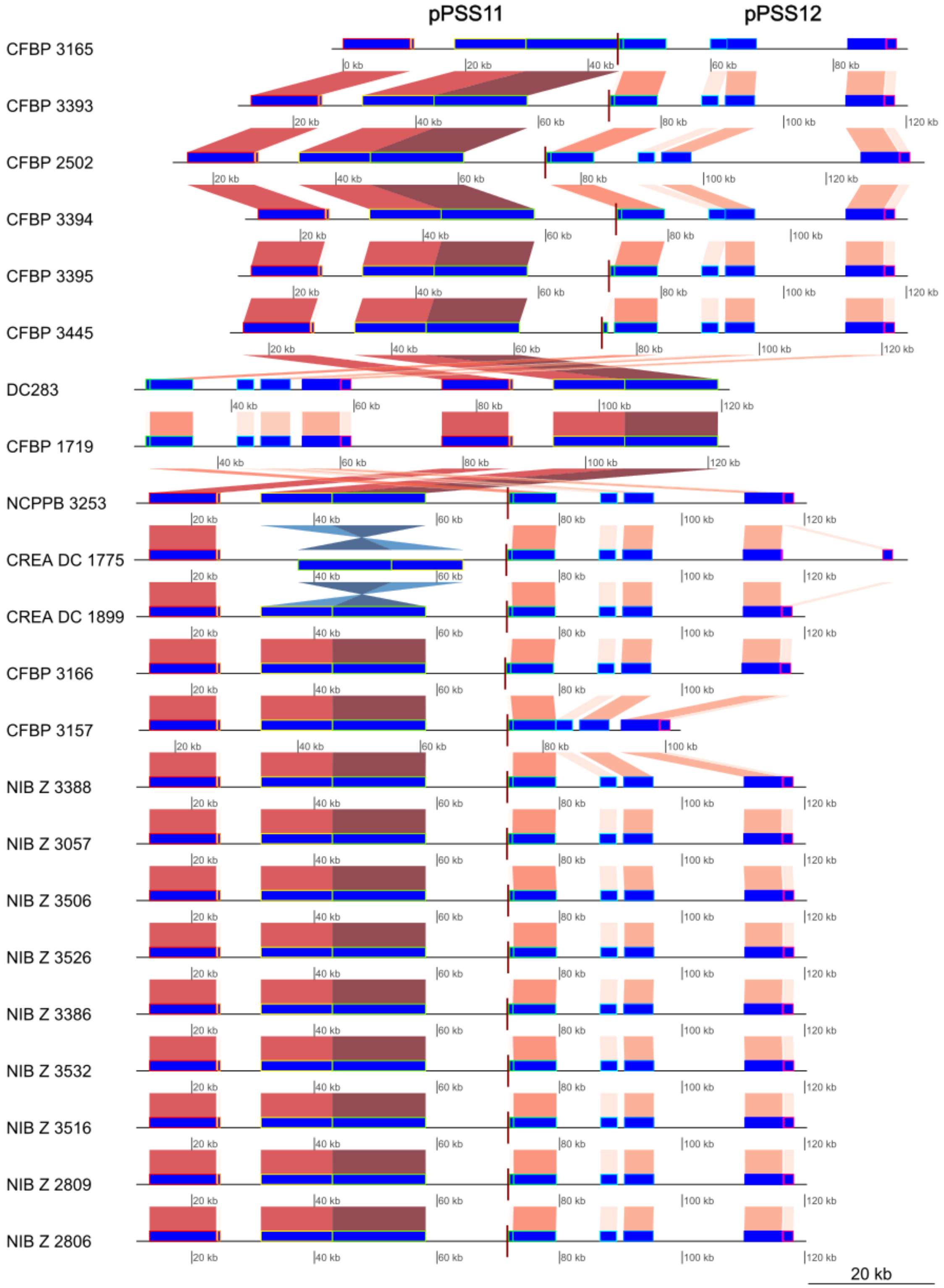

*Figure S2: Alignment showing all complete plasmids pPSS14 from assembled genomes of* Pantoea stewartii *subsp.* stewartii *strains in which this plasmid is present in complete form. The genomes of strains CFBP 3165 and CFBP 3169 lack a large segment of plasmid in comparison to other genomes. Strains are ordered based on their position in the phylogenetic tree in Figure 2A.*

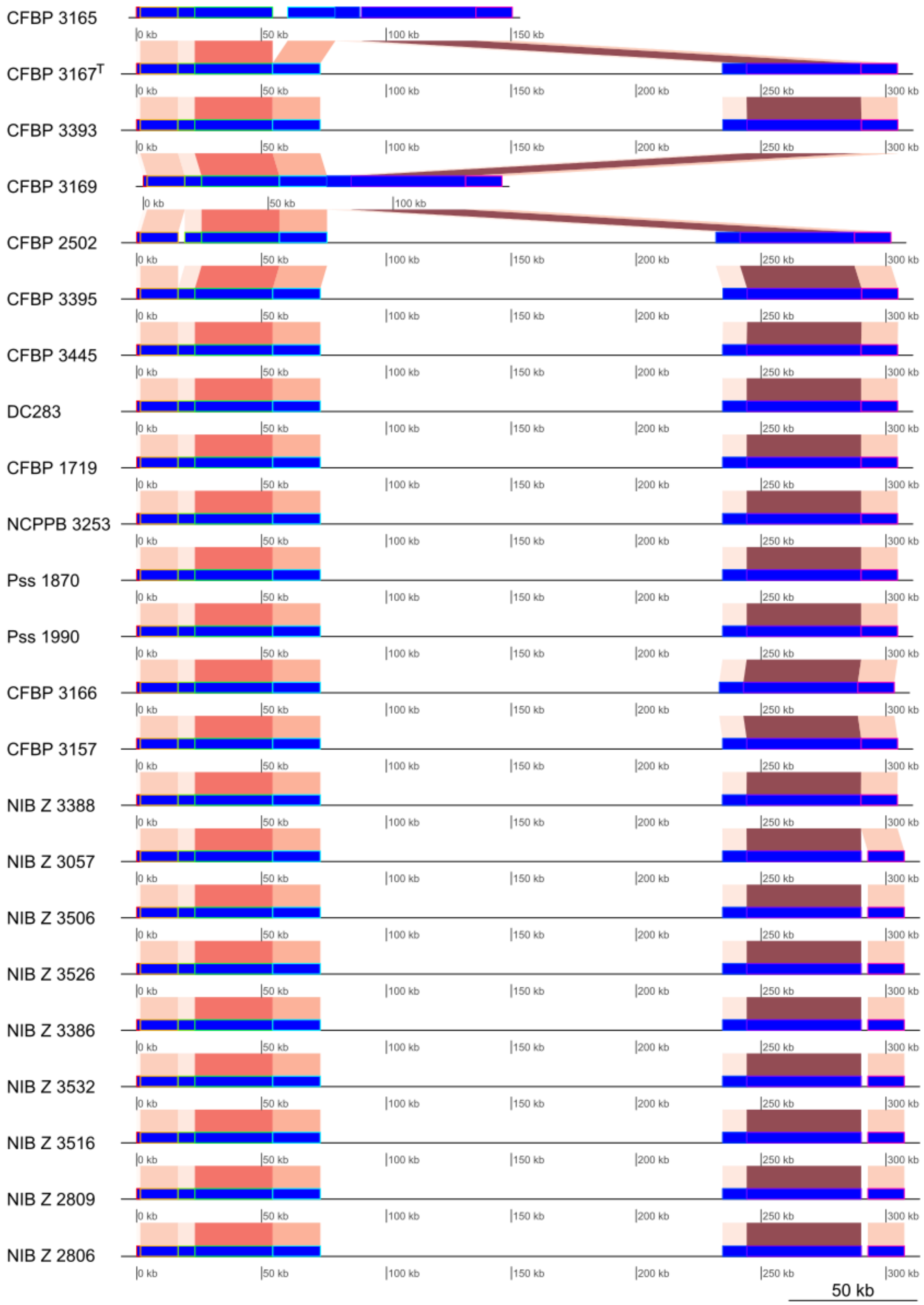

*Figure S3: Alignment showing all complete plasmids pPSS04, pPSS05, and pPSS07 from assembled genomes of* Pantoea stewartii *subsp.* stewartii *strains in which this plasmid is present in complete form. Assembled genomes contain either plasmid pPSS04 (mostly 25 kb) or plasmids pPSS05 (mostly 26 kb) and pPSS07 (mostly 47 kb). The exceptions are strains CFBP 3393, CFBP 2502, and CFBP 3394, which only contain plasmid pPSS05. In strain CFBP 3394, plasmid pPSS05 also contains an insertion from plasmid pPSS10 (Supplement Figure S7). Strains are ordered based on their position in the phylogenetic tree in Figure 2A.*

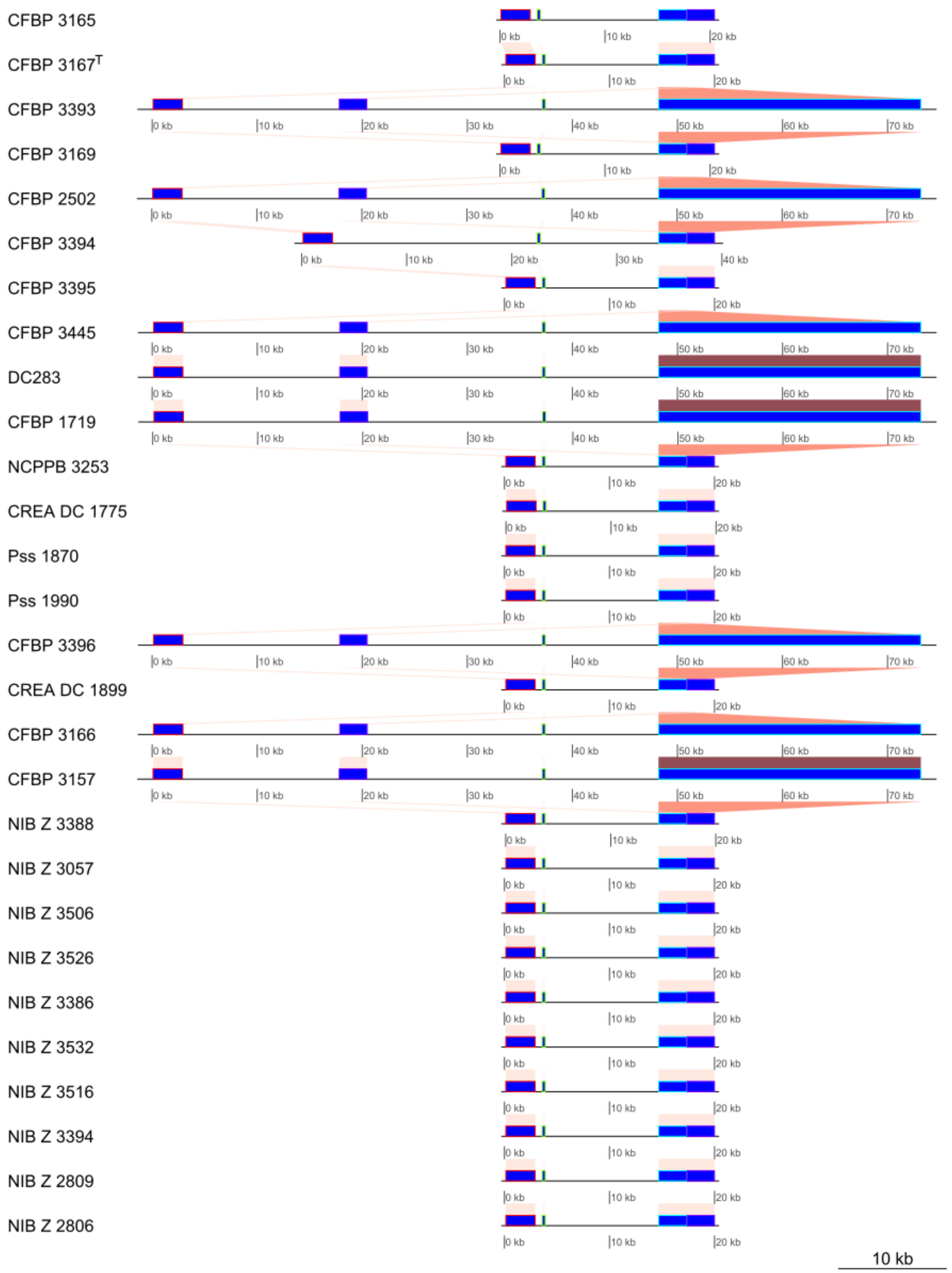

*Figure S4: Alignment showing all complete plasmid pPSS08 from assembled genomes of* Pantoea stewartii *subsp.* stewartii *strains in which this plasmid is present in complete form. This plasmid is present in genomes in either a small variant, which is mostly 34 kb in size, or a large variant, which is mostly 53 kb in size. Strains are ordered based on their position in the phylogenetic tree in Figure 2A.*

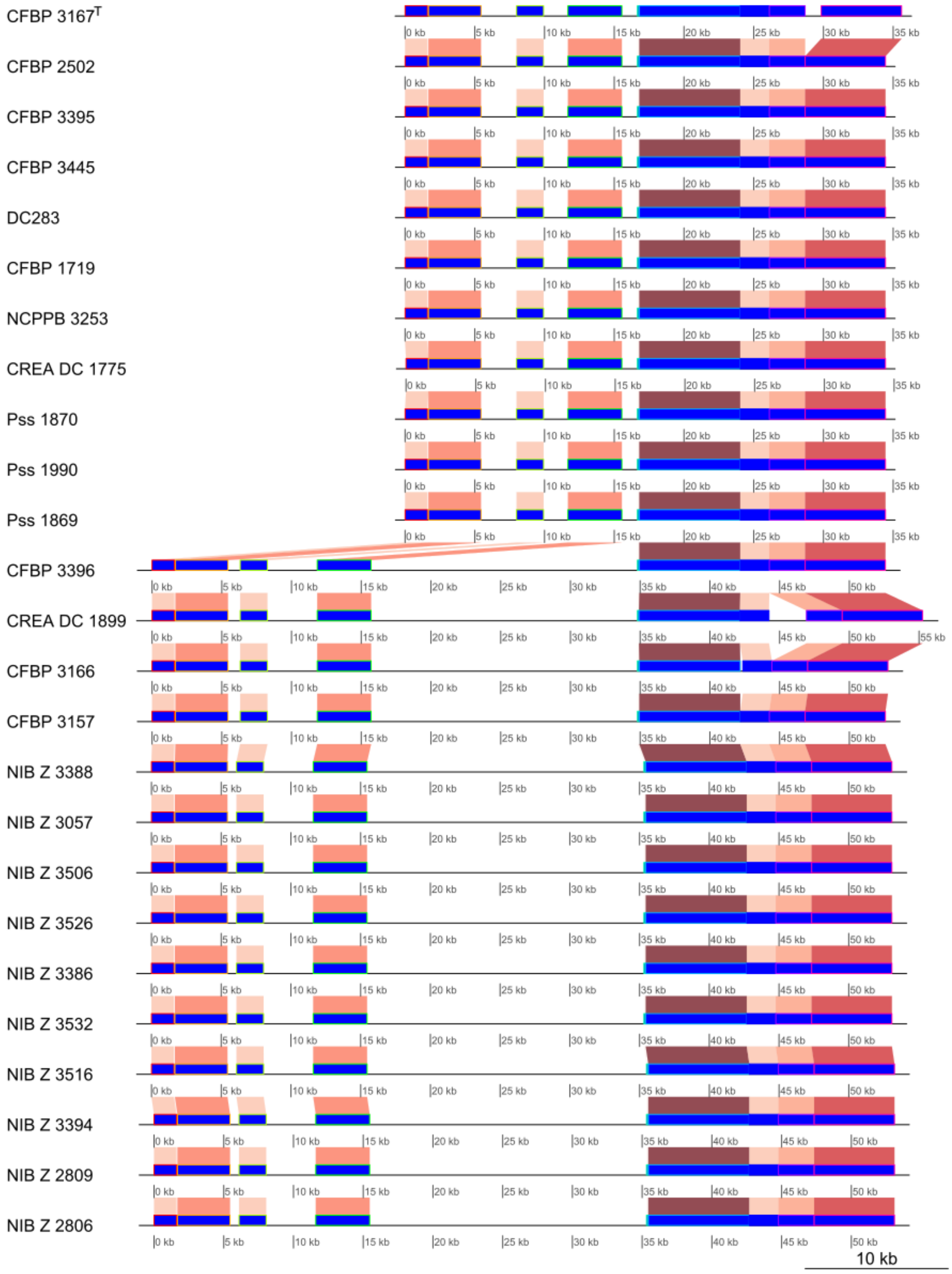

*Figure S5: Alignment showing plasmids pPSS03 and pPSS10 in genomes of* Pantoea stewartii *subsp.* stewartii *strains CFBP 3157 and DC283 (reference genome). In the genomes of strain CFBP 3157 segment of the plasmid pPSS10 is inserted into plasmid pPSS03.*

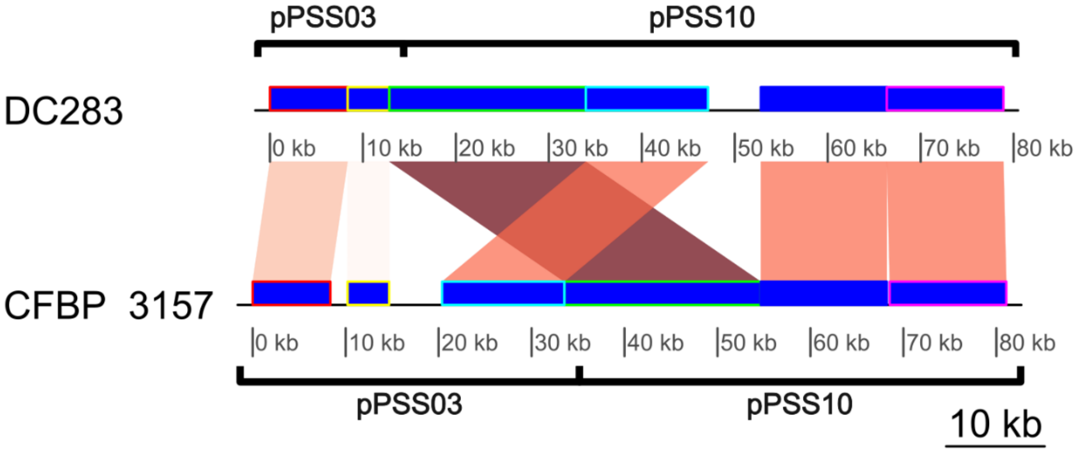

*Figure S6: Alignment showing plasmids pPSS05 and pPSS10 in genomes of* Pantoea stewartii *subsp.* stewartii *strains CFBP 3394 and DC283 (reference genome). In the genome of strain CFBP 3394 segment of the plasmid pPSS10 is inserted into plasmid pPSS05.*

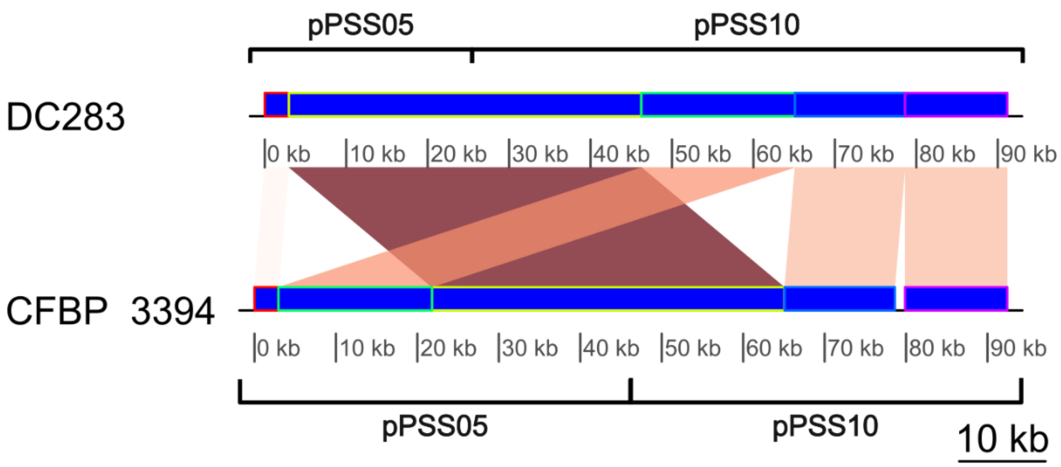
